# Advancing in silico drug design with Bayesian refinement of AlphaFold models

**DOI:** 10.1101/2025.06.25.661454

**Authors:** Samiran Sen, Samuel E. Hoff, Tatiana I. Morozova, Vincent Schnapka, Massimiliano Bonomi

**Affiliations:** Institut Pasteur, Université Paris Cité, CNRS UMR 3528, Computational Structural Biology Unit, Paris, France; CNRS, ENS de Lyon, LPENSL, UMR5672, 69342, Lyon cedex 07, France

## Abstract

Virtual screening has become an indispensable tool in modern structure-based drug discovery, enabling the identification of candidate molecules by computationally evaluating their potential to bind target proteins. The accuracy of such screenings critically depends on the quality of the target structures employed. Recent advances in protein structure prediction, particularly AlphaFold2, have revolutionized this field with unprecedented accuracy. However, AlphaFold2 models often exhibit limitations in local structural details, especially within binding pockets, which limit their utility for small molecule docking. In contrast, molecular dynamics simulations with accurate atomistic force fields can refine protein structures, but lack the ability to leverage the structural information provided by deep learning approaches. Here, we introduce bAIes, an integrative method that bridges this gap by combining physics-based force fields with data-driven predictions through Bayesian inference. Crucially, bAIes demonstrates a superior ability to discriminate between binders and non-binders in virtual screening campaigns, outperforming both AlphaFold2 and molecular dynamics-refined models. By enhancing the usability of AlphaFold2 models without requiring extensive experimental or computational resources, bAIes offers a convenient solution to a longstanding challenge in structure-based drug design, potentially accelerating the early phases of drug discovery.

## Introduction

Structure-based drug design (SBDD) is a cornerstone of modern-day drug discovery, integrating high-resolution structural data with advanced computational techniques. The foundation of this approach lies in determining how small molecules bind to target biomolecules at the atomic level. This enables the design of molecules with high affinity, selectivity, and specific pharmacological profiles, advancing drug development in areas like oncology, infectious diseases, and neurodegenerative disorders.^1^ In the context of SBDD, virtual screening (VS) has emerged as an efficient *in silico* approach to identify potential binders from large chemical libraries.^2–4^ By prioritizing candidate molecules for synthesis and testing, this procedure accelerates experimental validation and subsequent lead optimization. ^5^ The key components of this process are: *(i)* a method for obtaining accurate, ligand-receptive structural models of the target protein;^6,7^ *(ii)* a docking algorithm capable of efficiently exploring ligand binding poses;^8^ and *(iii)* an accurate scoring function to rank poses and compare ligands. ^9,10^

In this work, we focus on the generation of accurate models of the target. Since determining an experimental structure is often challenging and time consuming, researchers frequently rely on computational models. ^6,7^ Comparative modeling has traditionally been used when experimental structures of homologous proteins with high sequence identity are available.^8,11–13^ However, when the target lies in the twilight zone, *i*.*e*. with sequence identity below 30%, the reliability of comparative models drop drastically, and additional refinement becomes necessary. A potential solution to this problem came with the breakthrough in structure prediction by deep learning approaches such as AlphaFold2^14^ (AF), which opened up new possibilities for targeting systems lacking both experimental structures and homologs with high sequence identity. ^15^ With AF, the availability of high-accuracy protein models was extended to the entire human proteome,^14,16,17^ greatly expanding the scope of VS studies.

Using AF models in docking and VS studies, however, presents several challenges. One key issue is determining whether AF models, which are typically generated in absence of ligands, resemble more closely an *apo* (unbound) state, a *holo* (ligand-bound) state, or an intermediate between the two, as each conformation can have a distinct impact on docking performance. Although AF models generally display high structural accuracy, studies have shown that they often underperform compared to experimental *holo* structures when used as targets for docking.^18^ In some cases, AF models identify top-ranking ligands with a success rate comparable to that of experimental *apo* structures, but they still perform worse than *holo* conformations. ^19^ These findings suggest that AF models may be more *apo*-like in nature. However, in a few instances, AF models have successfully identified binders that experimental structures could not, ^20^ implying that they might occasionally capture different *holo*-like conformations. Finally, in cases where experimental structures are not available, comparative models still perform generally better than AF, except for targets in the twilight zone.^21^

In many of the instances mentioned above, the poor performance of AF models in docking and VS can be traced to inaccuracies in side chain conformations, such as steric clashes or unlikely rotamer states, ^22^ even when the backbone is predicted with high accuracy. Therefore, additional local refinement of AF models, particularly focused on side chain orientations, may be necessary to improve their utility and accuracy in SBDD. Several approaches have been developed to refine computational models with the goal of improving docking performance, with Molecular Dynamics (MD) simulations being one of the most popular approaches.^23–27^ These methods generally focus on local stereochemistry refinement through extensive MD simulations, typically with positional restraints on the initial model. ^28,29^ However, when these procedures rely solely on molecular-mechanics force fields, they do not consistently lead to improved docking outcomes,^30,31^ largely because of the limited accuracy of the force field when used without guidance from other sources of structural information, such as experimental data. ^32^

In this work, we introduce bAIes, a method that combines Alphafold2 predictions with state-of-the-art molecular mechanics force fields. At the core of this integration, bAIes quantifies the local uncertainties in an AF model and uses a Bayesian inference frame-work to optimally combine the structural information provided by this tool with accurate physico-chemical descriptions of the system. Here, we evaluate the performance of bAIes in identifying binders through screening of large small molecule libraries. bAIes consistently outperforms AF and MD models by ranking binders significantly higher than non-binders. Overall, we show that bAIes is able to provide a principled path toward making AF structural models truly actionable for drug discovery.

## Theory

In integrative structural biology, Bayesian modeling approaches are used to combine diverse experimental data with physico-chemical knowledge to generate accurate and precise structural models of biological systems. To optimally integrate different sources of information, these methods explicitly model uncertainties and errors, allowing each piece of data to be weighted according to its reliability. Similarly, bAIes models the uncertainty in structural information provided by AlphaFold2 (AF) using distograms, which are distance distributions between *C*_*β*_ atoms (or *C*_*α*_ atoms for glycines). Distograms measure the local confidence of the AF model and are used in the structure module to guide model generation. Consequently, the most probable *C*_*β*_–*C*_*β*_ distance in a distogram typically matches closely the corresponding distance in the final predicted structure.

Generally speaking, Bayesian inference methods estimate the probability of a model *M*, defined in terms of its structure **X** and other parameters, given the information available about the system, including prior physico-chemical knowledge and newly acquired data. The posterior probability *p*(*M* | *D*) of model *M* given data *D* and prior knowledge, is expressed as:

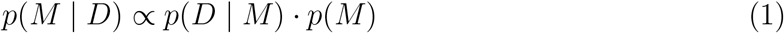

where the likelihood function *p*(*D* | *M*) represents the probability of observing data *D* given *M*. This function quantifies the agreement between the observed data and the model and accounts for all sources of uncertainty and error in the data. The term *p*(*M*) is the probability of model *M* based on prior knowledge. In the following, we specify each component of this general Bayesian framework as applied in our bAIes approach.

### The data

In integrative structural biology applications, *D* = {*d*_*i*_} represents a collection of experimental data points, often originating from different techniques, such as NMR, ^33^ SAXS,^34^ or cryo-electron microscopy.^35^ In bAIes, *D* represents the structural information provided by AF, defined as a collection of distances between *C*_*β*_ atoms (or *C*_*α*_ in the case of glycines). Each data point *d*_*i*_ represents the most probable distance computed from the corresponding AF distogram. Distances between residues separated in sequence by less than 3 aminoacids were discarded. Additionally, a residue-pair specific cutoff ^36^ was applied to retain only those residue pairs whose distances are indicative of potential inter-residue interactions.

### Data likelihood

For each pair of selected residues, a log-normal likelihood was used:

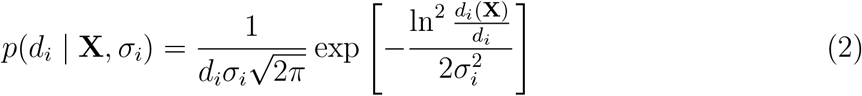

where *d*_*i*_(**X**) is the distance between a pair of residues in structure **X**, *d*_*i*_ the most probable distance in the corresponding AF distogram, and *σ*_*i*_ an error parameter that modulates the deviation between observed and computed data. Under the assumption of independent data points, the total likelihood for a set of of *m* points *D* = {*d*_*i*_} can be factorized as:

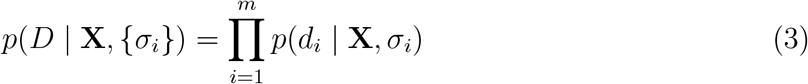

### Priors

In bAIes, the error parameters *σ*_*i*_ are treated as unknown variables and are determined by sampling the posterior distribution. To penalize large errors, we used an informative Jeffreys prior:^37^

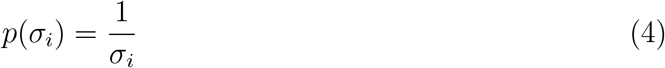

Additionally, each distogram was fit with a log-normal distribution whose standard deviation 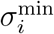 was used as minimum value for each uncertainty parameter. This choice allows us to handle cases where AF predictions are inaccurate in the same way we would treat outlier experimental data points in other Bayesian inference approaches for structural biology. Atomistic molecular mechanics force fields *E*_FF_ were used as structural prior:

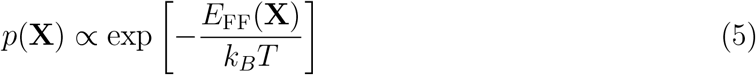

### Marginalization

As the product of each likelihood function with the corresponding uncertainty prior can be analytically integrated in *σ*_*i*_, to avoid explicit sampling of the uncertainty parameters we computed the marginal data likelihood as:^38^

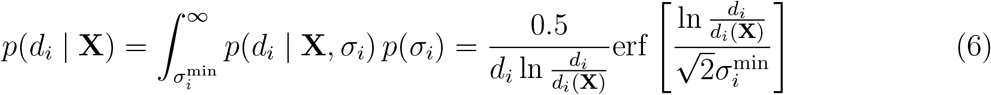

### The bAIes hybrid energy function

After defining all the components of our approach and marginalizing the uncertainty parameters, we obtain the final bAIes posterior distribution:

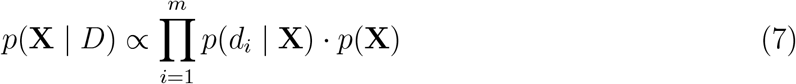

where the marginal data likelihood is given by Eq. 6 and the structural prior by Eq. 5. To sample the posterior, we define the associated bAIes hybrid energy function as:

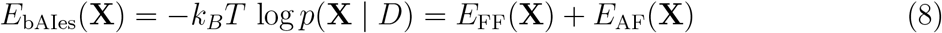

which is composed of the molecular mechanics force field *E*_FF_ and the spatial restraints *E*_AF_ that incorporate the structural information provided by the AF distograms. Finally, models are sampled by MD under the bAIes hybrid energy function defined in Eq. 8. In summary, during a bAIes simulation, the standard MD force field is modified by spatial restraints centered on the most probable distance predicted by AF and with intensities reflecting the AF confidence in the model (Fig. 1). Distograms with wide distributions result in weak structural restraints, while narrow distributions suggesting high confidence of AF in the model are translated into strong restraints.

**Figure 1.**
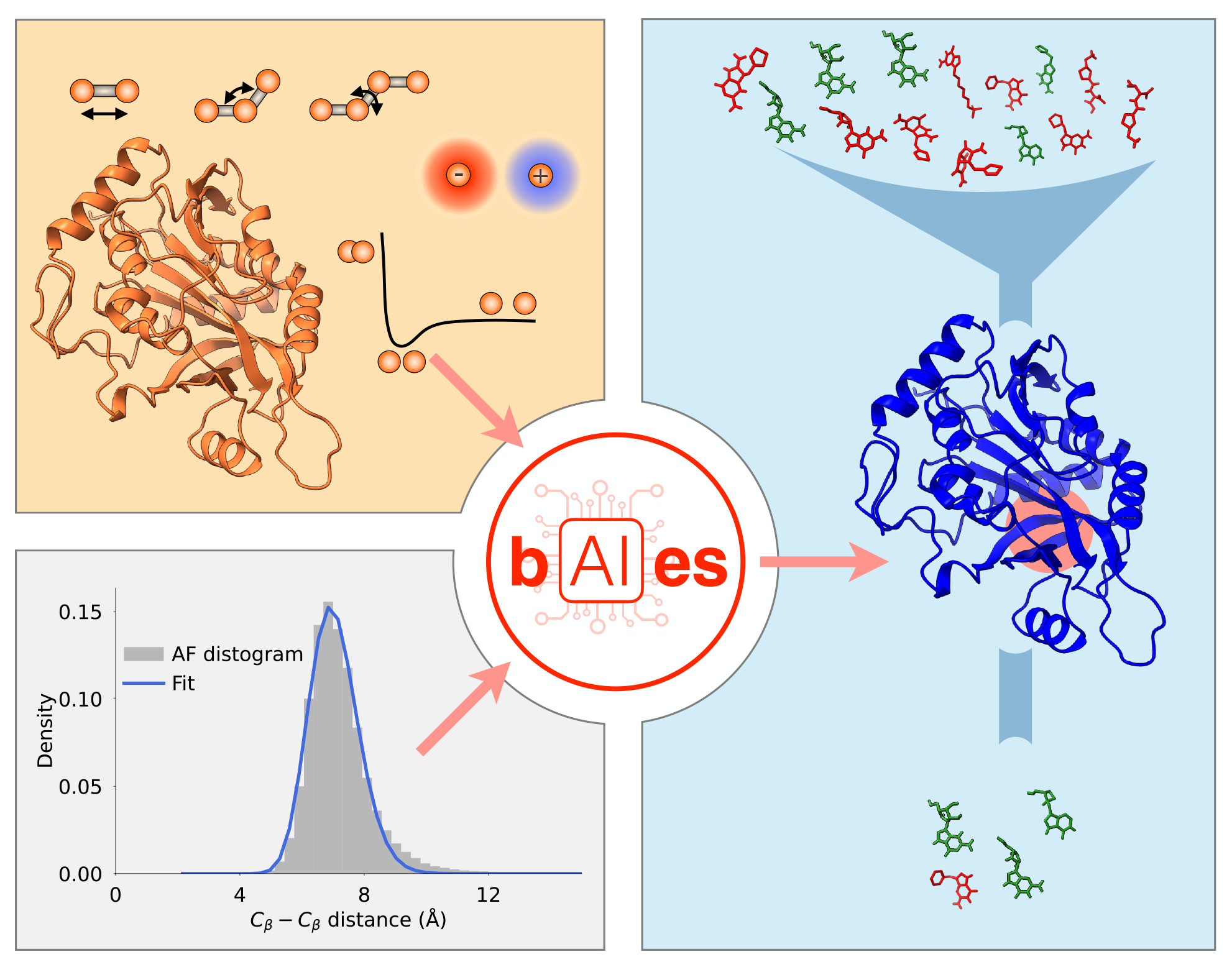
Overview of bAIes. bAIes is a Bayesian inference framework to integrate state-of-the-art molecular mechanics force fields (*top left*) with pairwise distograms reflecting the confidence in AlphaFold2 predictions (*bottom left*) to refine protein targets models and improve docking of large small molecule libraries (*right*).

## Results

This section is organized as follows. First, we analyze the bAIes-refined structures of ligand binding sites starting from AF models. We then evaluate whether bAIes models can be used in combination with docking protocols to *i)* reproduce the native ligand poses observed in experimental structures; and *ii)* discriminate between binders and non-binders during virtual screening of large libraries of small molecules (Fig. 1).

### Structural accuracy of ligand binding pockets

We evaluated the accuracy and precision of bAIes-refined ligand-binding pockets using a benchmark set of 15 proteins, whose structures in complex with their native ligands were determined by X-ray crystallography at high resolution (1.5 *−* 3.2Å, Table S1). This benchmark set is composed of systems of varying complexity, including soluble and transmembrane proteins, nuclear and cytoplasmic proteins, as well as monomeric and multimeric assemblies. To quantify model quality, we calculated the Root Mean Square Deviation (RMSD) of the *C*_*β*_ atoms of the ligand pocket from the corresponding experimental structure (Methods). The quality of bAIes-optimized pockets was compared to both the original AF models as well as those obtained by refining the AF models with MD simulations in presence (restrained) and absence (unrestrained) of harmonic positional restraints.

AF models were generally of good quality (mean pocket RMSD 1.0 Å) (Fig. S1, gray) and predicted with high confidence, consistent with prior reports. ^18,39^ We quantified the local level of confidence in the pocket region using the average pLDDT score across the binding pocket residues (binding pocket pLDDT). This was appreciably high, with values in most cases greater than 90, indicating that AF modeled these regions with high confidence. Notably, even the pockets that deviated significantly from the selected experimental *holo* structure were predicted with high pLDDT, suggesting that they might correspond to alternative conformational states. We investigated the conformations generated by unrestrained MD and restrained MD (rMD) by analyzing the distributions of pocket RMSDs along each trajectory. Unrestrained MD yielded pockets of broadly similar average accuracy to AF (mean pocket RMSD 1.53 Å). rMD offered a clear improvement over MD (mean pocket RMSD 1.10 Å). bAIes-refined pockets achieved a mean pocket RMSD of 1.16 Å, representing a similar improvement over MD and closely matching the accuracy of both AF and rMD.

Substantial structural variability of the binding pocket was also observed in all simulations, reflected in the width of the RMSD distribution within each trajectory. In the case of unrestrained MD, pocket RMSDs covered a broad range from 0.48Å (PDB 2AM9) to 2.02Å (PDB 3BGS) (Fig. S1, orange). As expected, in the case of rMD, we obtained narrower RMSD distributions, ranging from 0.15Å (PDB 1UDT and 2UZ3) to 0.31Å (PDB 3BGS) (Fig. S2, orange). In the case of bAIes, the RMSD distributions were similarly narrow, ranging from 0.21 Å (PDB 2UZ3) to 0.70 Å (PDB 2ICA) (Fig. S1, blue). In 67% of cases the sampled conformations included structures closer to the experimental reference than the AF model — compared to 33% of the cases for MD and 53% of the cases for rMD.

In the analysis reported in this section, we compared our models to a single experimental *holo* structure. However, it is possible that the original AF model, as well as the MD-, rMDand bAIes-refined conformations, resembled more closely a different *holo* structure. To test this hypothesis, we selected three systems (PDBs 3ERD, 3BGS, and 1H00) for which multiple *holo* structures in complex with different ligands were available (Table 1). We observed in each case that each of the AF, bAIes, MD, and rMD models resembled more closely a *holo* conformation different from the experimental reference initially selected (Fig. 2; Table 1, S2).

**Table 1.** Binding pocket structural accuracy. Binding pocket RMSD of the AF model and representatives of the top 3 clusters found by rMD and bAIes simulations with respect to the reference (*ref*.) as well as closest (*best*) experimental *holo* structures for the 3 systems used in the virtual screening. Also, the lowest RMSD obtained in each case

| System | PDB | AF model |  |  | rMD top 3 clusters |  |  | bAIs top 3 clusters |  |  |
| --- | --- | --- | --- | --- | --- | --- | --- | --- | --- | --- |
|  |  | RMSD | PDB | RMSD | RMSD | PDB | RMSD | RMSD | PDB | RMSD |
|  | ref. holo | ref. holo | best holo | best holo | ref. holo | best holo | best holo | ref. holo | best holo | best holo |
|  |  | [Å] |  | [Å] | [Å] |  | [Å] | [Å] |  | [Å] |
| Estrogen<br>receptor $\alpha$ | 3ERD | 0.85 | 1GWR | 0.39 | 0.91 | 1GWR | 0.46 | 0.92 | 1GWR | 0.46 |
|  |  |  |  |  | 0.91 | 1GWR | 0.48 | 0.97 | 1GWR | 0.45 |
|  |  |  |  |  | 0.89 | 1GWR | 0.50 | 1.00 | 1GWR | 0.49 |
| Purine<br>nucleoside<br>phosphorylase | 3BGS | 0.92 | 1RT9 | 0.79 | 1.21 | 1PF7 | 1.02 | 1.06 | 1RT9 | 0.92 |
|  |  |  |  |  | 1.19 | 2Q70 | 1.07 | 1.02 | 1RT9 | 0.89 |
|  |  |  |  |  | 1.10 | 2Q70 | 0.93 | 1.05 | 1RT9 | 0.92 |
| Cyclin-dependent<br>kinase | 1H00 | 5.37 | 3TI1 | 0.69 | 5.40 | 2VTN | 0.68 | 5.29 | 2VTN | 0.79 |
|  |  |  |  |  | 5.31 | 1WCC | 0.64 | 5.28 | 2VTN | 0.75 |
|  |  |  |  |  | 5.32 | 1WCC | 0.69 | 5.40 | 3TI1 | 0.90 |

**Figure 2.**
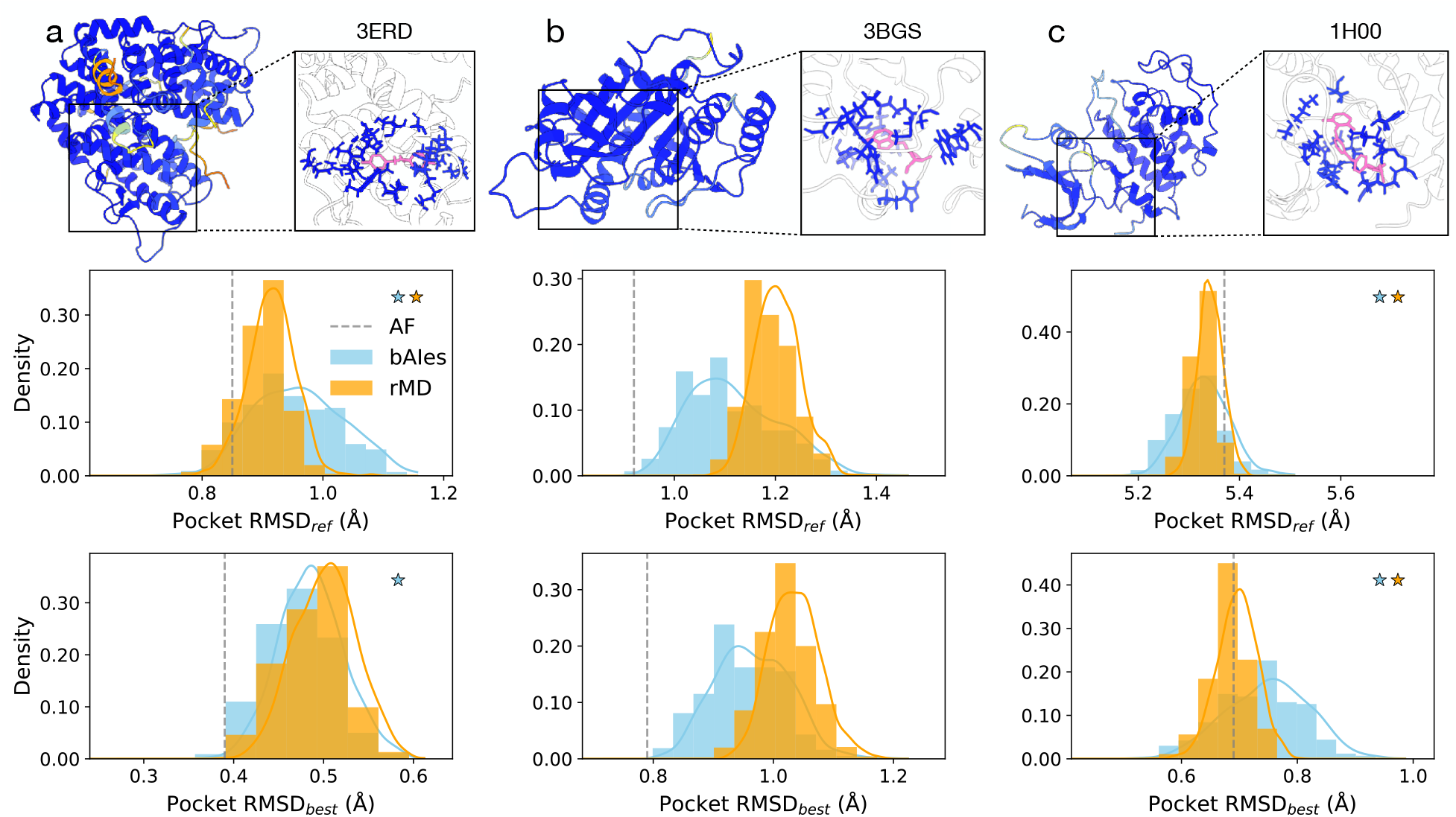
Structural accuracy of ligand binding pockets. Assessment of structural accuracy of ligand binding pockets for 3 selected systems: (**a**) Estrogen receptor *α* (PDB 3ERD), (**b**) Purine nucleoside phosphorylase (PDB 3BGS), and (**c**) Cyclin-dependent Kinase (PDB 1H00). For each system: (*top*) AF model of the protein, colored according to pLDDT, with inset highlighting the binding site residues (pLDDT-colored, licorice) and the ligand positioned as in the corresponding X-ray structure (pink, licorice); (*middle*) distributions of binding pocket *C*_*β*_-RMSD from the reference experimental *holo* conformation, for bAIes (blue) and rMD (orange) simulations. RMSD of the AF model is represented with a gray dashed line; (*bottom*) distributions of binding pocket *C*_*β*_-RMSD calculated from the closest experimental *holo* conformation. Orange and blue stars mark whether rMD and bAIes, respectively, sample pocket conformations closer to the experimental one than AF. Raw normalized histograms are represented by bars, kernel density estimations as solid lines.

### Structural accuracy of native ligand docking poses

Here we evaluate the impact of target structure refinement on docking accuracy. To do this, we docked the (native) ligands from our benchmark set of 15 systems (Table S1) to the original AF model as well as to selected bAIes-, MD-, and rMD-refined conformations. Typically, multiple docking poses (up to 9) were generated, usually differing in energy values of ~ 1kcal mol^*−*1^ between the highest and lowest scored models. Docking success was measured by computing the minimum ligand RMSD (RMSD_*l*_) between the reference experimental *holo* structure and the top five poses ranked by the docking scoring function (Methods). A docking trial was considered successful when the RMSD_*l*_ was below 2Å.

Re-docking the native ligands to their corresponding experimental *holo* structures was consistently successful (Fig. 3a, pink). All systems yielded RMSD_*l*_ below 2Å, with 33% achieving values below 1Å, confirming the validity of our docking protocol ^40^ (Fig. 3b, pink). In contrast, docking native ligands to the AF models was successful in only 40% of cases (Fig. 3a, gray), with RMSD_*l*_ ranging from 0.81 to 1.72Å and an average of 1.39Å (Fig. 3b, gray).

**Figure 3.**
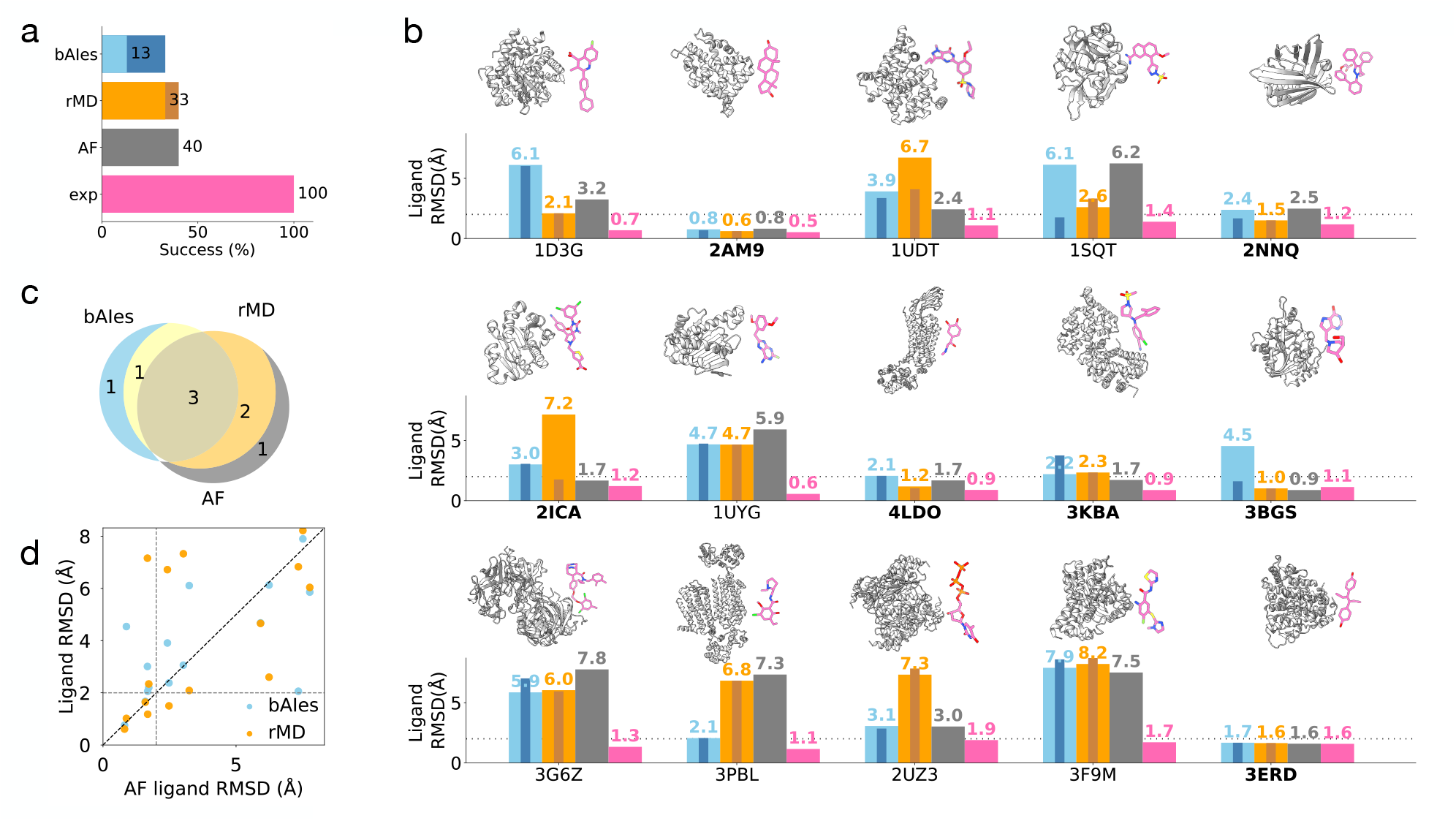
Structural accuracy of native ligand docking poses. **(a)** Percentage of docking successes across the entire benchmark set of 15 systems using the top1 cluster for bAIes (blue) and rMD (orange) as well as using AF (gray) and experimental (pink) structures. Also, the percentage of successes when top3 clusters are included for bAIes (dark blue), and rMD (dark orange). **(b)** For each system in the benchmark: (*top*) cartoon representation of the experimental *holo* structure, with highlight on the native ligand (pink); (*bottom*) ligand RMSDs with respect to experimental structure obtained from docking against top1 clusters of bAIes and rMD, as well as AF, and experimental structures. Bar coloring as in **(a).** Also, RMSDs when top3 clusters are included for bAIes (inner dark blue bar) and rMD (inner dark orange bar). Systems that resulted in a successful dock by either bAIes, rMD or AF models are marked with the PDB name in bold. **(c)** Diagram illustrating the overlap among successful cases of docking by bAIes, rMD, and AF models. Numbers represent unique and shared successes among the models; coloring as in **(a). (d)** Correlation between ligand RMSDs when docking against AF models and either bAIes (blue dots) or rMD models (orange dots). The ligand RMSDs values corresponding to the cutoff for successful docking (2Å) are indicated by gray dashed lines.

For MD, rMD, and bAIes models, each trajectory was first clustered based on structural similarity, and the centers of the top 3 populated clusters were selected as docking targets (Methods). Finally, to evaluate docking accuracy, the ligand pose with minimum ligand RMSD among the top five ranked poses was selected. Accuracy was evaluated on poses obtained by docking only against the most populated cluster (top1) or across the three most populated clusters (top3). In the case of MD, successful docking was observed in 13% of the systems (Fig S4a, orange) when the top1 or top3 clusters were considered. In the latter case, the RMSD_*l*_ ranged from 0.64 to 1.58Å, with an average of 1.11Å (Fig. S3, S4b, orange). These results were comparable with those obtained with the AF models. In the case of rMD, successful docking was observed in 33% and 40% of the systems (Fig. 3a, orange) when the top1 and top3 clusters were considered, respectively. In the latter case, the RMSD_*l*_ ranged from 0.61 to 1.75Å, with an average of 1.26Å, also comparable with the results obtained with AF models (Fig. 3a, orange; Fig. S3). For bAIes-refined models, successful docking was observed in 13% and 33% of the systems when the top1 and top3 clusters were considered, respectively (Fig. 3a, blue). In the successful cases obtained with the top3 clusters, the RMSD_*l*_ ranged from 0.68 to 1.75Å, with an average of 1.48Å, also closely matching the performance of AF models (Fig. 3b, blue; Fig. S3). In these subset of systems, bAIes accuracy was higher than MD and rMD in 80% and 20% of cases, respectively (Fig. 3b,d).

Overall, we employed 3 distinctly different ways to model protein structures: deep-learning prediction (AF), molecular dynamics (MD), and Bayesian refinement combining AF and MD (bAIes). We wanted to explore the possibility of combining these methods or a subset of them to maximize the chances of success in docking native ligands. We first noticed that MD (or rMD) docking was never successful in cases where either AF or bAIes failed (Fig. 3c, Fig. S4). Furthermore, in cases where either AF or bAIes was successful, these two methods produced the most accurate pose in the majority of cases. Based on these considerations, we tested a combination of only AF and bAIes as follows. We therefore pooled the docking poses from both AF and bAIes models and re-ranked them together. This approach yielded success rates similar to those of the individual methods alone, suggesting that combining data from bAIes and AF docking does not provide any particular advantage. This somewhat disappointing result can largely be attributed to the limitations of the scoring function, which was not always capable of ranking the most accurate poses highly.

### Virtual screening of small molecule libraries

In real-life drug discovery campaigns, docking is primarily used to screen large libraries of small molecules against protein targets, often with unknown *holo* structures, to identify potential binders. For such tasks, the ability to reliably discriminate binders from non-binders is often more critical than achieving highly accurate docking poses, as the ultimate goal is to prioritize candidates for experimental validation rather than to precisely model binding interactions. In this section, we assess the ability of bAIes to discriminate between binders and non-binders. To do so, we selected three systems for which large ligand libraries containing both binders and decoys were available (Methods) and that exhibited mixed native-ligand docking performance across the different models (Table S3). For each system, we screened the libraries against AF, MD, rMD, and bAIes models, as well as one experimental *holo* structure. The three selected targets were the Estrogen receptor *α* (PDB 3ERD), the Purine nucleoside phosphorylase (PDB 3BGS), and the Cyclin-dependent Kinase (PDB 1H00), which were screened against a total of 58409 small molecules, of which 1422 were known binders (Fig. 4a).

**Figure 4.**
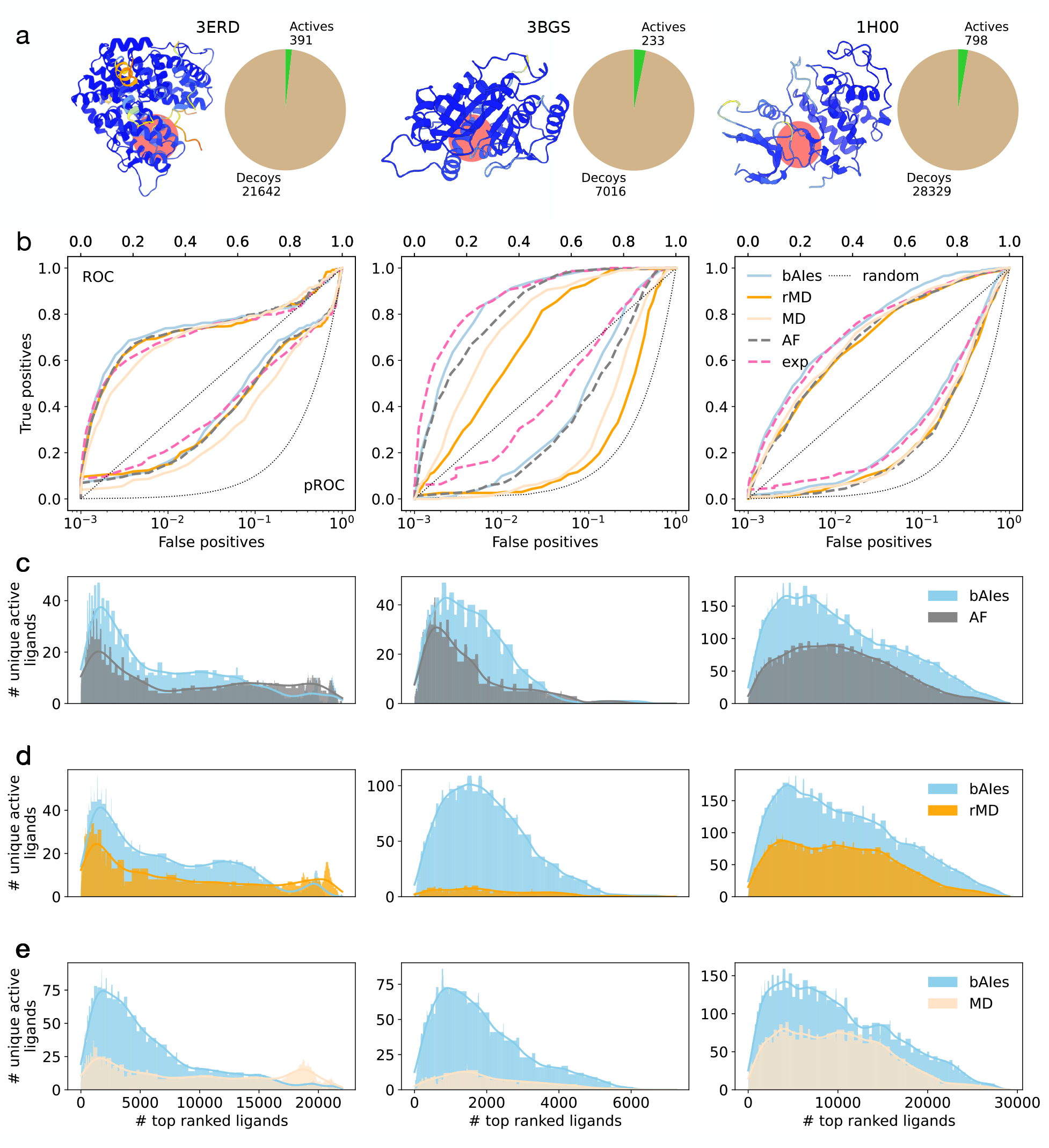
Virtual screening of small molecule libraries. Assessment of the ability of different models to discriminate between actives and decoys in large-scale small molecule libraries, for 3 selected systems: (*left*) Estrogen receptor *α* (PDB 3ERD), (*center*) Purine nucleoside phosphorylase (PDB 3BGS), and (*right*) Cyclin-dependent Kinase (PDB 1H00). (**a**) AF model of the protein target (pLDDT-colored), with highlighted binding pocket (coral-colored disc) alongside a pie chart with the number of active (green) and decoy (beige) ligands present in the library. (**b**) ROC (upper diagonal) and pROC (lower diagonal) curves obtained with the bAIes (blue), rMD (orange), MD (light orange), AF (gray dashed) models, and experimental structures (pink dashed). Also, the random classifier for ROC and pROC (black dotted). (**c-e**) Number of unique active ligands identified by (**c**) bAIes and AF models, bAIes and rMD models, and (**e**) bAIes a2n6d MD models, as a function of the number of top-ranked ligands; kernel density estimations (solid lines colors as in (**b**)).

In order to quantify the performance of each method used to model the target, we used two metrics: the receiver operating characteristic (ROC) curve^41^ and the linear-log ROC (pROC) curve. ^42^ Briefly, the ROC curve characterizes how well a method can distinguish between true positives (actual binders) and false positives (non-binders) as the score threshold (or rank) for selecting molecules is varied. The area under the ROC curve (ROC-AUC) provides a single measure of this performance: a larger AUC indicates a better ability to rank true binders ahead of non-binders, with an AUC of 1.0 representing perfect classification and 0.5 representing random guessing. The pROC curve extends this idea by placing greater emphasis on the early part of the ranking, which is particularly important in virtual screening contexts where only the top-ranked compounds are typically tested. ^43,44^ Therefore, a larger area under the pROC curve (pROC-AUC) reflects a method’s effectiveness in identifying true binders among the top-scoring candidates.

First, we observed that, for all the three targets studied, both the experimental structure (Fig. 4b, pink) as well as the AF (Fig. 4b, gray), MD (Fig. 4b, light orange), rMD (Fig. 4b, orange), and bAIes (Fig. 4b, blue) models had a ROC-AUC larger than a random classifier (Fig. 4b, dotted gray). Furthermore, the experimental structures showed excellent discriminatory power, with average ROC-AUC and pROC-AUC of 0.79 and 1.24, respectively. The AF models also performed well (average ROC-AUC of 0.76), but showed early enrichment consistently lower than that obtained using the experimental structure (average pROC-AUC of 1.03). Notably, in the case of the estrogen receptor *α*, the AF ROC-AUC of 0.74 was slightly superior to the experimental one of 0.73, though the early enrichment was still poorer (pROC-AUC of 1.21 compared to 1.28).

In the case of MD, rMD, and bAIes, the target models were selected by clustering the trajectories in the same way as for native ligand docking (Methods). Due to the large computational cost, only the center of the top cluster was used for virtual screening. The ROC-AUCs obtained with the MD model were consistently worse than the experimental ones by 2.7%, 13.5% and 4.1% for PDB 3ERD, 3BGS, 1H00, respectively. The pROC-AUCs were also significantly lower than the experimental ones by 19.5%, 45.6% and 16.5% for PDB 3ERD, 3BGS, and 1H00, respectively (Table 2). When comparing with AF, the performance varied across systems. The MD ROC-AUCs were worse for PDB 3ERD and 3BGS (by 4.1% and 8.3%, respectively) and so were the corresponding pROC-AUCs (by 14.9% and 25.9%, respectively). For the third system (PDB 1H00), the MD ROC-AUC and pROC-AUC were slightly better than AF (by 1.4% and 2.5%, respectively) (Table 2).

**Table 2.** Virtual screening performances with experimental (exp), AF, MD, rMD and bAIes structures. Area under the ROC curve (ROC-AUC) and pROC curve (pROC-AUC) for the 3 systems used in the virtual screening.

|  | 3ERD |  | 3BGS |  | 1H00 |  |
| --- | --- | --- | --- | --- | --- | --- |
|  | ROC-AUC | pROC-AUC | ROC-AUC | pROC-AUC | ROC-AUC | pROC-AUC |
| exp | 0.73 | 1.28 | 0.89 | 1.47 | 0.74 | 0.97 |
| AF | 0.74 | 1.21 | 0.84 | 1.08 | 0.7 | 0.79 |
| MD | 0.71 | 1.03 | 0.77 | 0.8 | 0.71 | 0.81 |
| rMD | 0.73 | 1.19 | 0.70 | 0.69 | 0.70 | 0.7 |
| bAIs | 0.75 | 1.23 | 0.86 | 1.16 | 0.75 | 0.94 |

In the case of rMD, the ROC-AUCs were worse than (21.3% and 5.7% for PDB 3BGS and 1H00, respectively) or comparable to the experimental ones (PDB 3ERD). The pROC-AUCs were also significantly worse in all cases (7.0%, 53.1% and 27.8% for PDB 3ERD, 3BGS, and 1H00, respectively; Table 2). When comparing to the AF models, the performance was poorer in two cases: the ROC-AUCs were worse by 1.4% and 16.7%, and the pROC-AUCs were worse by 1.7% and 36.1%, for PDB 3ERD and 3BGS, respectively. In contrast with MD, in the case of the third system (PDB 1H00), the rMD ROC-AUC was comparable to the AF model, while the pROC-AUC was still worse (by 11.4%) (Table 2). Overall, using rMD models in VS did not provide a clear advantage over MD.

Remarkably, bAIes models had a higher enrichment than the experimental structures for two of our targets (by 2.7% and 1.4% for PDB 3ERD and 1H00, respectively), but worse in the other case (by 3.4% for PDB 3BGS). However, pROC-AUCs were consistently worse than the experimental ones by 3.9%, 21.1% and 3.1% for PDB 3ERD, 3BGS, and 1H00, respectively (Table 2). In comparison with the AF model, bAIes consistently showed better discriminatory performance across all systems studied (ROC-AUC higher by 1.4%, 2.4% and 7.1% for PDB 3ERD, 3BGS, and 1H00, respectively). Notably, the early enrichments were significantly better, with pROC-AUC higher by 1.7%, 7.4% and 19.0% for PDB 3ERD, 3BGS, and 1H00, respectively (Table 2).

When compared to MD, bAIes consistently produced significantly higher ROC-AUC (by 5.6%, 11.7% and 5.6% for PDB 3ERD, 3BGS, and 1H00, respectively) as well as superior pROC-AUC (by 19.4%, 45.0% and 16.0% for PDB 3ERD, 3BGS and 1H00, respectively). When compared to rMD, bAIes also consistently produced significantly higher ROC-AUC (by 2.7%, 22.9% and 7.1% for PDB 3ERD, 3BGS, and 1H00, respectively), and also superior pROC-AUC (by 33.6%, 68.1% and 34.3% for PDB 3ERD, 3BGS, and 1H00, respectively). Notably, the bAIes improvement in early enrichment with respect to MD, rMD and AF is 2 to 5 times the improvement observed in the overall discriminative capacity. This indicates that, in absence of available experimental structures, bAIes is the best single approach for obtaining early enrichment compared to MD, rMD, and AF models.

Although AF, MD, and rMD showed inferior performance in early enrichment, it is worth exploring whether the distinct conformations generated by these methods enabled the identification of some specific binders that bAIes failed to capture. To investigate this, we calculated the number of unique ligands identified by each method across the three systems studied, as a function of the ranking selection threshold. We found that the AF model was able to capture, among the top ranked ligands, a significant number of binders not identified by bAIes in two out of three cases (PDB 3ERD and 3BGS) (Fig. 4c); in the third case (PDB 1H00) bAIes clearly outperformed AF. We also noticed that between the rMD and MD models, there was no clear advantage of one over the other: the rMD models identified significantly greater, fewer and equal number of top-ranked unique binders in the case of PDB 3ERD, 3BGS, and 1H00, respectively (Fig. S5). On the other hand, bAIes consistently identified a significantly higher number of unique ligands compared to rMD (Fig. 4d) and MD (Fig. 4e). These observations suggest that, when sufficient computational resources are available, using both AF and bAIes models as target structures in virtual screening campaigns may be a valuable strategy to identify a diverse set of ligands for experimental testing.

## Discussion

In this work, we introduce bAIes, a Bayesian modeling framework that leverages the structural information provided by AF to refine atomistic protein models for use in virtual screening of small molecules. bAIes combines the structural information encoded in AF distograms with state-of-the-art molecular mechanics force fields employed in molecular dynamics simulations (Theory). Notably, across our virtual-screening benchmarks, bAIes-refined models consistently outperformed both the original AF predictions and structures refined by standard MD, both in presence and absence of positional restraints. In particular, our models demonstrated a superior ability to distinguish binders from non-binders, especially among the top-scoring ligands. This enhanced early enrichment is critical in real-world virtual screening campaigns, where only the highest-ranked compounds are typically selected for experimental validation.

Our results suggest that bAIes is able to sample conformations that are close to the initial AF models, representative of local structural fluctuations of the target, and suitable for ligand binding. In principle, such conformations should also be accessible through standard MD simulations. However, limitations in the underlying MD force fields may hinder the sampling of these conformations. By integrating the pairwise distance distributions provided by AF, we were able to overcome these limitations. An intriguing interpretation of these findings is that distograms provide not only a measure of confidence in the AF predictions, but also encode information about protein dynamics. This idea is supported by our recent work in which we used bAIes to determine accurate atomistic structural ensembles of intrinsically disordered proteins.^45^ Additionally, we demonstrated that enriching graph-based protein representations with distograms consistently improves performance across diverse protein and RNA property prediction tasks, even outperforming methods that rely on computationally expensive MD simulations.^46^

We also evaluated the ability of bAIes to refine ligand-binding pockets. In this context, we obtained mixed results. The bAIes models from our benchmark of 15 systems did not demonstrate a clear advantage over directly using the AF-generated structures. However, it remains to be systematically assessed whether AF and bAIes captured *holo*-like (or *apo*) conformations that differ from the one that we designated as reference structure. This would explain the performances observed both when assessing the accuracy of the ligand-binding pockets and with the native ligand docking. The analysis of three systems in our benchmark set seemed to support this hypothesis (Table 1, Fig. 2).

To overcome some of the limitations described above, several potential improvements to bAIes can be envisioned. First, in this work, we performed only short simulations to refine the models. In the future, thanks to the integration of bAIes into the open-source PLUMED library,^47,48^ it will be straightforward to combine our method with enhanced sampling techniques, such as metadynamics,^47^ to access conformations further distant from the initial AF models. Second, in this study, conformations for docking or virtual screening were selected solely based on cluster populations. Moving forward, conformations generated by bAIes could be re-scored using scoring functions and consensus strategies specifically designed to identify druggable conformations.^49–51^ Finally, docking protocols and scoring functions play a critical role in correctly ranking compounds. We foresee a more systematic evaluation of bAIes performances using a broader range of docking approaches, particularly new methods based on deep learning.^52,53^

In this work, we used distograms generated by AlphaFold2, as the source code for AlphaFold3 (AF3) was not available at the time we developed our approach, and distogram information could not be retrieved from the AF3 web server. Now that the source code has been released, it is possible to obtain AF3 distograms, which could further expand the range of applications for bAIes. For example, bAIes could be employed to refine RNA models for use in virtual screening. Targeting RNAs with small molecules is an exciting and rapidly growing field of research,^54,55^ attracting increasing interest from pharmaceutical companies. Moreover, AF3 now allows for modeling protein and RNA targets in complex with small molecules. The poses obtained with AF3 could be further refined with bAIes by integrating distograms that describe target-ligand distance distributions. It would be interesting to assess whether bAIes-refined models of small molecule–protein complexes improve ligand pose accuracy or virtual screening enrichment. Nevertheless, limitations in the usage and licensing of AF3 outputs may complicate further development of bAIes based on this new version of the software.

In conclusion, bAIes provides a computationally efficient approach to refine AlphaFold2 models of protein targets for their use in the early stages of drug design. The method is implemented in the PLUMED-ISDB module^56^ of the open-source PLUMED library (www.plumed.org),^47,48^ making it readily accessible to a large computational science community. Our approach has the potential to improve the reliability of virtual screening campaigns by prioritizing compounds with a higher likelihood of success. As such, it offers a valuable addition to modern drug discovery pipelines, where the availability of accurate target models remains a key bottleneck. Ultimately, bAIes can contribute to the development of more effective and selective therapeutic candidates.

## Methods

### AlphaFold2 model generation

To generate the AF models, the complete FASTA sequence of each target protein (Table S1) was fed into AlphaFold2 v2.3.2. ^14^ The multimer model was used, and MSA searches were conducted with HHBlits ^57^ and HHSearch^58^ on BFD^14^ and Uniref30 (until 30-02-2023) databases, and with Jackhmmer from HMMER3^59^ on Uniref90^60^ and clustered MGnify ^61^ databases. All genetic databases (full databases) used at CASP14 were searched for optimizing the structures. Default templates (until 14-05-2020) were used to guide the structure predictions. A single seed per model was used to make 5 predictions that were ranked according to their model confidences, measured by the predicted LDDT (pLDDT). The top ranked model was relaxed with gradient descent by OpenMM using the AMBER99SB^62^ force field and selected for further analysis.

### Molecular dynamics and bAIes simulations

All simulations were performed with GROMACS v2022.5^63^ equipped with PLUMED v2.9. ^47^

#### Setup and equilibration

For soluble proteins, the AF models were solvated in a cubic box, with dimensions chosen to ensure that its boundaries were at least 1.0nm away from any modeled atom. Na^+^ and Cl^*−*^ ions were added at a concentration of 0.15M to neutralize the net system charge. The AMBER99SB-ILDN^64^ force field and TIP3P model^65^ were used for protein and water molecules, respectively. Equations of motion were integrated with a standard leap-frog algorithm with a timestep of 1fs. Van der Waals interactions were gradually switched off at 0.8nm and cut off at 0.9nm; the fast smooth particle-mesh Ewald (PME) method was used to calculate electrostatic interactions with cutoff at 0.9nm. ^66^ Energy-minimization was performed on the solvated structures with the steepest-descent algorithm. This was followed by relaxation first under NVT conditions for 100ps to a target temperature of 300K. The Bussi-Donadio-Parrinello thermostat ^67^ was used with a time constant of 0.2ps. The systems were then equilibrated in the NPT ensemble for 100ps to a target pressure of 1bar using an isotropic Berendsen barostat^68^ with coupling time constant of 1ps. During both NVT and NPT equilibration, harmonic restraints were applied to the positions of the protein heavy atoms, with force constant equal to 4000kJ mol^*−*1^ nm^*−*2^.

For membrane proteins (PDB 4LDO and 3PBL), the AF models were first oriented in an implicit lipid bilayer parallel to the *xy* plane using the PPM 3.0 webserver,^69^ and then used as input for CHARMM-GUI.^70^ The systems were inserted in a homogeneous POPC lipid bilayer and solvated in a triclinic box. Na^+^ and Cl^*−*^ ions were added at a concentration of 0.15M to neutralize the net system charge. The CHARMM36m^71^ and mTIP3P model^72^ were used for protein/lipids and water molecules, respectively. Van der Waals interactions were gradually switched off at 1.0nm and cut off at 1.2nm; PME was used to calculate electrostatic interactions with cutoff at 1.2nm. ^66^ The standard CHARMM-GUI protocol was used to equilibrate the systems. Multiple consecutive short simulations in the NVT and NPT ensembles at a temperature of 303.15K and pressure of 1bar were performed to equilibrate the systems. The Bussi-Donadio-Parrinello thermostat ^67^ was used with a time constant of 0.2ps. For the NPT runs, a semi-isotropic Berendsen barostat ^68^ was used to allow deformations in the *xy* plane independent from the *z*-axis. During these equilibration steps, harmonic restraints on the positions of the lipid and protein atoms were gradually switched off. Equations of motion were integrated with a standard leap-frog algorithm with a timestep of 1fs or 2fs, depending on the stage of equilibration.

#### Production

MD, restrained MD (rMD), and bAIes structural refinement production simulations were run for 5ns post-equilibration. For soluble proteins, simulations were performed in the NVT ensemble with a timestep of 1fs. Temperature was set at 300K using the Bussi-Donadio-Parrinello thermostat with time constant of 0.2ps. For membrane proteins, simulations were performed in the NPT ensemble with a timestep of 2fs. Temperature was set at 303.15K using the Bussi-Donadio-Parrinello as for soluble proteins; pressure was set at 1bar using the semi-isotropic Parrinello-Rahman barostat ^73^ with time constant of 5ps. For rMD simulations, harmonic restraints were applied to the positions of the protein heavy atoms, with force constant equal to 4000kJ mol^*−*1^ nm^*−*2^. For bAIes simulations, restraints were applied with frequency equal to 4fs.

#### Analysis

To select representative models from the MD, rMD and bAIes trajectories, all generated conformations were clustered according to their binding pocket structure as follows. First, we defined the pocket as the set of protein residues comprising heavy atoms within 6Å of any heavy atom of the ligand in the X-ray structure. Then we employed the *gromos* clustering approach ^74^ using the RMSD calculated on the *C*_*β*_ pocket atoms as metrics. Several clustering cutoffs were tested ranging from 0.01 to 0.13nm. A system-specific cutoff was chosen to ensure a balance between having clusters that were conformationally homogeneous within and distinct from each other.

### Benchmark of structural accuracy of ligand binding pockets

#### Details of the systems

15 systems were chosen to benchmark the accuracy of different methods to refine the target structure. The *holo* conformations of these systems were determined by X-ray crystallography at resolution ranging from 1.5Å to 3.2Å (Table S1). The systems included a mixture of soluble and membrane proteins. The soluble proteins were: Dihydroorotate dehydrogenase (PDB 1D3G), Androgen Receptor (PDB 2AM9), Phosphodiesterase 5A (PDB 1UDT), Urokinase-type plasminogen activator (PDB 1SQT), Fatty-acid-binding protein 4 (PDB 2NNQ), Leukocyte-function associated antigen 1 (PDB 2ICA), Heat shock protein 90 *α* (PDB 1UYG), Progesterone receptor (PDB 3KBA), Purine nucleoside phosphorylase (PDB 3BGS), Renin (PDB 3G6Z), Thymidine kinase (PDB 2UZ3), Hexokinase type IV (PDB 3F9M), and Estrogen receptor *α* (PDB 3ERD). The membrane proteins were: *β*_2_ adrenergic receptor (PDB 4LDO) and Dopamine *D*_3_ receptor (PDB 3PBL). Each target structure was resolved in complex with a native ligand (Table S1). Among these systems, 9 were monomers (PDBs 1D3G, 2AM9, 1UDT, 1SQT, 2NNQ, 2ICA, 1UYG, 3BGS, and 3F9M), 3 were homodimers (PDBs 3KBA, 3G6Z, and 3PBL), 1 was a homotetramer (PDB 2UZ3), 1 was a heterodimer (PDB 4LDO), and 1 was a heterotetramer (PDB 3ERD).

#### Analysis

In order to compare the different methods (AF, MD, rMD, and bAIes) used in this work, each protein model was first aligned to the corresponding experimental structure. Due to possible missing residues in the experimental structure, we first performed a sequence alignment to the model sequence using Biopython. ^75^ We then used the sequence alignment to perform structural alignment of the AF model, and every frame of the MD, rMD, and bAIes trajectories, to the reference experimental structure using MDAnalysis v2.7.0. ^76,77^ The structural alignments and RMSD calculations were performed in the same way as described for the clustering of the trajectories.

### Benchmark of structural accuracy of native ligand docking poses

#### Details of the systems

The same 15 systems defined in the previous section were used to benchmark the accuracy of native ligand docking.

#### Details of the docking protocol

Models produced by the different methods (AF, MD, rMD, and bAIes) were first stripped of any lipids, ions and water molecules present, and empirical atomic partial charges in the form of Gasteiger charges were added. The experimental structures were prepared for docking by removing the native ligands present in it. Autodock Vina 1.1.2^78,79^ was used to dock each ligand to its receptor. A moderate exhaustiveness of 8 was used to specify the number of independent runs starting from random ligand conformations for each docking trial. This value is typically considered a good trade-off between computational cost and accuracies in Autodock Vina. ^80^ A cubic box centered at the centre of mass of the experimental native ligand structure and with side equal to 20Å was used for docking. The default united-atom scoring function was used to rank the docked ligands. ^78^

#### Analysis

For each docked ligand, the ligand RMSD (RMSD_*l*_) with respect to the experimental structure was calculated. In the case of MD, rMD, and bAIes, the structure(s) used for docking was chosen in two different ways: (i) the center of the most populated cluster (top1), and (ii) the collection of centers of three most populated clusters (top3). Thereafter, in each of the two cases, the 5 top scoring poses were considered. Out of these 5 poses, the one with lowest RMSD_*l*_ was selected as the representative ligand pose. A docking trial was considered successful if RMSD_*l*_ *<* 2Å.

### Virtual screening of small molecule libraries

#### Details of the systems

The following 3 systems were chosen for benchmarking the virtual screening of large libraries of small molecules against the AF, MD, rMD, and bAIes models: Estrogen receptor *α* (PDB 3ERD), Purine nucleoside phosphorylase (PDB 3BGS), and the Cyclin-dependent Kinase (PDB 1H00). The structure of these systems were determined by X-ray crystallography at high resolution (PDB 3ERD: 2.0Å; PDB 3BGS: 2.1Å; PDB 1H00: 1.6Å). Two of these systems were monomers (PDB 3BGS and 1H00) and one was a heterotetramer (PDB 3ERD).

#### Details of the small molecule libraries

For the estrogen receptor *α*, a library of 391 actives and 21642 decoys was downloaded from the DUD-E database.^81^ For the purine nucleoside phosphorylase, a library of 233 actives and 7016 decoys was downloaded from NRLiSt BDB. ^82^ For the cyclin-dependent kinase, a library of 798 actives and 28329 decoys was downloaded from the DUD-E database.

#### Details of the virtual screening protocol

For each system of our benchmark, the AF model, the top1 MD, rMD, and bAIes clusters, and the experimental structure were prepared for virtual screening using the same protocol defined for native ligand docking.

#### Analysis

To analyze the output of our virtual screening, only the top scoring pose of each small molecule was considered. The docking performance of each method was then evaluated independently for each system in the benchmark set in the following way. First, we sorted the small molecules by their docking scores, then we computed the ROC curve by counting the number of true and false positives as the docking score threshold to select molecules was varied. We measured early enrichment quality of the receptors with the pROC curve, for which we used a log_10_ scale for the false positives (Fig. 4b).

## Software and data availability

bAIes is implemented as a part of the PLUMED-ISDB module^56^ in the development version (GitHub master branch) of PLUMED (https://github.com/plumed/plumed2).^47^ The GROMACS topologies and PLUMED input files used in our benchmarks are available in PLUMED-NEST (www.plumed-nest.org), the public repository of the PLUMED consortium,^48^ as plumID:25.016. Scripts to prepare bAIes simulations as well as a complete tutorial are available on PLUMED-TUTORIALS (www.plumed-tutorials.org) ^83^ as ID:25.002.

## Acknowledgments

We thank Quang Tru Huynh for technical support with the Maestro cluster at Institut Pasteur. This project has received funding from the European Research Council (ERC) under the European Union’s Horizon 2020 research and innovation programme (Grant agreement No. 101086685 – bAIes).

## References

(1) DiMasi, J. A.; Hansen, R. W.; Grabowski, H. G. The price of innovation: new estimates of drug development costs. Journal of Health Economics 2003, 22, 151–185.

(2) Villoutreix, B. O.; Eudes, R.; Miteva, M. A. Structure-based virtual ligand screening: recent success stories. Combinatorial chemistry & high throughput screening 2009, 12, 1000–1016.

(3) Irwin, J. J.; Shoichet, B. K.; Mysinger, M. M.; Huang, N.; Colizzi, F.; Wassam, P.; Cao, Y. Automated Docking Screens: A Feasibility Study. Journal of Medicinal Chemistry 2009, 52, 5712–5720.

(4) Lavecchia A. D. G. C., Virtual Screening Strategies in Drug Discovery: A Critical Review. Current Medicinal Chemistry 2013, 20, 2839–2860.

(5) Walters, W.; Stahl, M. T.; Murcko, M. A. Virtual screening—an overview. Drug Discovery Today 1998, 3, 160–178.

(6) Shoichet, B. K. Virtual screening of chemical libraries. Nature 2004, 432, 862–865.

(7) Beddell, C.; Goodford, P.; Norrington, F.; Wilkinson, S.; Wootton, R. Compounds designed to fit a site of known structure in human haemoglobin. British journal of pharmacology 1976, 57, 201–209.

(8) Tuccinardi, T. Docking-based virtual screening: recent developments. Combinatorial chemistry & high throughput screening 2009, 12, 303–314.

(9) Brooijmans, N.; Kuntz, I. D. Molecular recognition and docking algorithms. Annual review of biophysics and biomolecular structure 2003, 32, 335–373.

(10) Halperin, I.; Ma, B.; Wolfson, H.; Nussinov, R. Principles of docking: An overview of search algorithms and a guide to scoring functions. Proteins: Structure, Function, and Bioinformatics 2002, 47, 409–443.

(11) Bordogna, A.; Pandini, A.; Bonati, L. Predicting the accuracy of protein–ligand docking on homology models. Journal of computational chemistry 2011, 32, 81–98.

(12) Cavasotto, C. N.; Phatak, S. S. Homology modeling in drug discovery: current trends and applications. Drug discovery today 2009, 14, 676–683.

(13) Holcomb, M.; Chang, Y.-T.; Goodsell, D. S.; Forli, S. Evaluation of AlphaFold2 structures as docking targets. Protein Science 2023, 32, e4530.

(14) Jumper, J.; Evans, R.; Pritzel, A.; Green, T.; Figurnov, M.; Ronneberger, O.; Tunyasuvunakool, K.; Bates, R.; Zídek, A.; Potapenko, A.; others Highly accurate protein structure prediction with AlphaFold. Nature 2021, 596, 583–589.

(15) Hameduh, T.; Haddad, Y.; Adam, V.; Heger, Z. Homology modeling in the time of collective and artificial intelligence. Computational and Structural Biotechnology Journal 2020, 18, 3494–3506.

(16) Tunyasuvunakool, K. et al. Highly accurate protein structure prediction for the human proteome. Nature 2021, 596, 590–596.

(17) Porta-Pardo, E.; Ruiz-Serra, V.; Valentini, S.; Valencia, A. The structural coverage of the human proteome before and after AlphaFold. PLOS Computational Biology 2022, 18, 1–17.

(18) Scardino, V.; Di Filippo, J. I.; Cavasotto, C. N. How good are AlphaFold models for docking-based virtual screening? iScience 2023, 26, 105920.

(19) Zhang, Y.; Vass, M.; Shi, D.; Abualrous, E.; Chambers, J. M.; Chopra, N.; Higgs, C.; Kasavajhala, K.; Li, H.; Nandekar, P.; Sato, H.; Miller, E. B.; Repasky, M. P.; Jerome, S. V. Benchmarking Refined and Unrefined AlphaFold2 Structures for Hit Discovery. Journal of Chemical Information and Modeling 2023, 63, 1656–1667.

(20) Lyu, J. et al. AlphaFold2 structures guide prospective ligand discovery. Science 2024, 384, eadn6354.

(21) Kersten, C.; Clower, S.; Barthels, F. Hic Sunt Dracones: Molecular Docking in Uncharted Territories with Structures from AlphaFold2 and RoseTTAfold. Journal of Chemical Information and Modeling 2023, 63, 2218–2225.

(22) Ruff, K. M.; Pappu, R. V. AlphaFold and Implications for Intrinsically Disordered Proteins. Journal of Molecular Biology 2021, 433, 167208.

(23) Feig, M. Local Protein Structure Refinement via Molecular Dynamics Simulations with locPREFMD. Journal of Chemical Information and Modeling 2016, 56, 1304–1312.

(24) Miller, E. B. et al. Reliable and Accurate Solution to the Induced Fit Docking Problem for Protein–Ligand Binding. Journal of Chemical Theory and Computation 2021, 17, 2630–2639.

(25) Wang, J.; Koirala, K.; Do, H. N.; Miao, Y. PepBinding: A Workflow for Predicting Peptide Binding Structures by Combining Peptide Docking and Peptide Gaussian Accelerated Molecular Dynamics Simulations. The Journal of Physical Chemistry B 2024, 128, 7332–7340.

(26) Chipot, C. Recent Advances in Simulation Software and Force Fields: Their Importance in Theoretical and Computational Chemistry and Biophysics. The Journal of Physical Chemistry B 2024, 128, 12023–12026.

(27) Bhattacharya, D.; Cheng, J. 3Drefine: Consistent protein structure refinement by optimizing hydrogen bonding network and atomic-level energy minimization. Proteins: Structure, Function, and Bioinformatics 2013, 81, 119–131.

(28) Heo, L.; Feig, M. What makes it difficult to refine protein models further via molecular dynamics simulations? Proteins: Structure, Function, and Bioinformatics 2018, 86, 177–188.

(29) Heo, L.; Janson, G.; Feig, M. Physics-based protein structure refinement in the era of artificial intelligence. Proteins: Structure, Function, and Bioinformatics 2021, 89, 1870–1887.

(30) Feig, M.; Mirjalili, V. Protein structure refinement via molecular-dynamics simulations: what works and what does not? Proteins: Structure, Function, and Bioinformatics 2016, 84, 282–292.

(31) Feig, M. Computational protein structure refinement: almost there, yet still so far to go. Wiley Interdisciplinary Reviews: Computational Molecular Science 2017, 7, e1307.

(32) Bonomi, M.; Heller, G. T.; Camilloni, C.; Vendruscolo, M. Principles of protein structural ensemble determination. Current Opinion in Structural Biology 2017, 42, 106– 116, Epub 2017 Jan 5.

(33) Heller, G. T. et al. Small-molecule sequestration of amyloid-β as a drug discovery strategy for Alzheimer’s disease. Science Advances 2020, 6, eabb5924.

(34) Jussupow, A.; Messias, A. C.; Stehle, R.; Geerlof, A.; Solbak, S. M.; Paissoni, C.; Bach, A.; Sattler, M.; Camilloni, C. The dynamics of linear polyubiquitin. Science Advances 2020, 6, eabc3786.

(35) Hoff, S. E.; Thomasen, F. E.; Lindorff-Larsen, K.; Bonomi, M. Accurate model and ensemble refinement using cryo-electron microscopy maps and Bayesian inference. PLOS Computational Biology 2024, 20, 1–26.

(36) Kamisetty, H.; Ovchinnikov, S.; Baker, D. Assessing the utility of coevolution-based residue–residue contact predictions in a sequence- and structure-rich era. Proceedings of the National Academy of Sciences 2013, 110, 15674–15679.

(37) Sivia, D.; Skilling, J. Data Analysis: A Bayesian Tutorial, 2nd ed.; Oxford University Press: Oxford, 2006.

(38) Bonomi, M.; Hanot, S.; Greenberg, C. H.; Sali, A.; Nilges, M.; Vendruscolo, M.; Pellarin, R. Bayesian Weighing of Electron Cryo-Microscopy Data for Integrative Structural Modeling. Structure 2019, 27, 175–188.e6.

(39) Karelina, M.; Noh, J. J.; Dror, R. O. How accurately can one predict drug binding modes using AlphaFold models? eLife 2023, 12, RP89386.

(40) Gohlke, H.; Hendlich, M.; Klebe, G. Knowledge-based scoring function to predict protein-ligand interactions11Edited by R. Huber. Journal of Molecular Biology 2000, 295, 337–356.

(41) Zweig, M. H.; Campbell, G. Receiver-operating characteristic (ROC) plots: a fundamental evaluation tool in clinical medicine. Clinical chemistry 1993, 39, 561–577.

(42) Knight, I. S.; Naprienko, S.; Irwin, J. J. Enrichment Score: a better quantitative metric for evaluating the enrichment capacity of molecular docking models. arXiv 2023, 2210.10905.

(43) Wei, B. Q.; Baase, W. A.; Weaver, L. H.; Matthews, B. W.; Shoichet, B. K. A Model Binding Site for Testing Scoring Functions in Molecular Docking. Journal of Molecular Biology 2002, 322, 339–355.

(44) Zhao, W.; Hevener, K.; White, S.; Lee, R.; Boyett, J. A statistical framework to evaluate virtual screening. BMC bioinformatics 2009, 10, 225.

(45) Schnapka, V.; Morozova, T. I.; Sen, S.; Bonomi, M. Atomic resolution ensembles of intrinsically disordered proteins with Alphafold. Nature Communications 2026, 17, 2399.

(46) Portal, N.; Karroucha, W.; Mallet, V.; Bonomi, M. Learning Dynamic Protein Representations at Scale with Distograms. bioRxiv 2026, 2026.01.29.702509.

(47) Tribello, G. A.; Bonomi, M.; Branduardi, D.; Camilloni, C.; Bussi, G. PLUMED 2: New feathers for an old bird. Computer Physics Communications 2014, 185, 604–613.

(48) PLUMED consortium Promoting transparency and reproducibility in enhanced molecular simulations. Nature Methods 2019, 16, 670–673.

(49) Sarkar, A.; Concilio, S.; Sessa, L.; Marrafino, F.; Piotto, S. Advancements and novel approaches in modified AutoDock Vina algorithms for enhanced molecular docking. Results in Chemistry 2024, 7, 101319.

(50) Singh, N.; Chaput, L.; Villoutreix, B. O. Fast Rescoring Protocols to Improve the Performance of Structure-Based Virtual Screening Performed on Protein–Protein Interfaces. Journal of Chemical Information and Modeling 2020, 60, 3910–3934.

(51) Graves, A. P.; Shivakumar, D. M.; Boyce, S. E.; Jacobson, M. P.; Case, D. A.; Shoichet, B. K. Rescoring docking hit lists for model cavity sites: predictions and experimental testing. Journal of Molecular Biology 2008, 377, 914–934.

(52) Gentile, F.; Agrawal, V.; Hsing, M.; Ton, A.-T.; Ban, F.; Norinder, U.; Gleave, M. E.; Cherkasov, A. Deep Docking: A Deep Learning Platform for Augmentation of Structure Based Drug Discovery. ACS Central Science 2020, 6, 939–949.

(53) Zheng, L.; Meng, J.; Jiang, K.; Lan, H.; Wang, Z.; Lin, M.; Li, W.; Guo, H.; Wei, Y.; Mu, Y. Improving protein–ligand docking and screening accuracies by incorporating a scoring function correction term. Briefings in Bioinformatics 2022, 23, bbac051.

(54) Panei, F. P.; Torchet, R.; Ménager, H.; Gkeka, P.; Bonomi, M. HARIBOSS: a curated database of RNA-small molecules structures to aid rational drug design. Bioinformatics 2022, 38, 4185–4193.

(55) Panei, F. P.; Gkeka, P.; Bonomi, M. Identifying small-molecules binding sites in RNA conformational ensembles with SHAMAN. Nature Communications 2024, 15, 5725.

(56) Bonomi, M.; Camilloni, C. Integrative structural and dynamical biology with PLUMED-ISDB. Bioinformatics 2017, 33, 3999–4000.

(57) Johnson, L. S.; Eddy, S. R.; Portugaly, E. Hidden Markov model speed heuristic and iterative HMM search procedure. BMC bioinformatics 2010, 11, 1–8.

(58) Steinegger, M.; Meier, M.; Mirdita, M.; Vöhringer, H.; Haunsberger, S. J.; Söding, J. HH-suite3 for fast remote homology detection and deep protein annotation. BMC bioinformatics 2019, 20, 1–15.

(59) Eddy, S. R. Accelerated profile HMM searches. PLoS computational biology 2011, 7, e1002195.

(60) Suzek, B. E.; Wang, Y.; Huang, H.; McGarvey, P. B.; Wu, C. H.; Consortium, U. UniRef clusters: a comprehensive and scalable alternative for improving sequence similarity searches. Bioinformatics 2015, 31, 926–932.

(61) Mitchell, A. L.; Almeida, A.; Beracochea, M.; Boland, M.; Burgin, J.; Cochrane, G.; Crusoe, M. R.; Kale, V.; Potter, S. C.; Richardson, L. J.; others MGnify: the microbiome analysis resource in 2020. Nucleic acids research 2020, 48, D570–D578.

(62) Hornak, V.; Abel, R.; Okur, A.; Strockbine, B.; Roitberg, A.; Simmerling, C. Comparison of multiple Amber force fields and development of improved protein backbone parameters. Proteins: Structure, Function, and Bioinformatics 2006, 65, 712–725.

(63) Abraham, M.; Murtola, T.; Schulz, R.; Páll, S.; Smith, J.; Hess, B.; Lindahl, E. GROMACS: High performance molecular simulations through multi-level parallelism from laptops to supercomputers. SoftwareX 2015, 1, 19–25.

(64) Lindorff-Larsen, K.; Piana, S.; Palmo, K.; Maragakis, P.; Klepeis, J. L.; Dror, R. O.; Shaw, D. E. Improved side-chain torsion potentials for the Amber ff99SB protein force field. Proteins: Structure, Function, and Bioinformatics 2010, 78, 1950–1958.

(65) Jorgensen, W. L.; Chandrasekhar, J.; Madura, J. D.; Impey, R. W.; Klein, M. L. Comparison of simple potential functions for simulating liquid water. The Journal of Chemical Physics 1983, 79, 926–935.

(66) Essmann, U.; Perera, L.; Berkowitz, M. L.; Darden, T.; Lee, H.; Pedersen, L. G. A smooth particle mesh Ewald method. The Journal of Chemical Physics 1995, 103, 8577–8593.

(67) Bussi, G.; Donadio, D.; Parrinello, M. Canonical sampling through velocity rescaling. The Journal of Chemical Physics 2007, 126, 014101.

(68) Berendsen, H. J.; Postma, J. v.; Van Gunsteren, W. F.; DiNola, A.; Haak, J. R. Molecular dynamics with coupling to an external bath. The Journal of chemical physics 1984, 81, 3684–3690.

(69) Lomize, A. L.; Todd, S. C.; Pogozheva, I. D. Spatial arrangement of proteins in planar and curved membranes by PPM 3.0. Protein Science 2022, 31, 209–220.

(70) Jo, S.; Kim, T.; Iyer, V. G.; Im, W. CHARMM-GUI: a web-based graphical user interface for CHARMM. Journal of computational chemistry 2008, 29, 1859–1865.

(71) Huang, J.; Rauscher, S.; Nawrocki, G.; Ran, T.; Feig, M.; De Groot, B. L.; Grubmüller, H.; MacKerell Jr, A. D. CHARMM36m: an improved force field for folded and intrinsically disordered proteins. Nature methods 2017, 14, 71–73.

(72) MacKerell Jr, A. D.; Bashford, D.; Bellott, M.; Dunbrack Jr, R. L.; Evanseck, J. D.; Field, M. J.; Fischer, S.; Gao, J.; Guo, H.; Ha, S.; others All-atom empirical potential for molecular modeling and dynamics studies of proteins. The journal of physical chemistry B 1998, 102, 3586–3616.

(73) Parrinello, M.; Rahman, A. Polymorphic transitions in single crystals: A new molecular dynamics method. Journal of Applied Physics 1981, 52, 7182–7190.

(74) Daura, X.; Gademann, K.; Jaun, B.; Seebach, D.; van Gunsteren, W. F.; Mark, A. E. Peptide Folding: When Simulation Meets Experiment. Angewandte Chemie International Edition 1999, 38, 236–240.

(75) Cock, P. J. A.; Antao, T.; Chang, J. T.; Chapman, B. A.; Cox, C. J.; Dalke, A.; Friedberg, I.; Hamelryck, T.; Kauff, F.; Wilczynski, B.; de Hoon, M. J. L. Biopython: freely available Python tools for computational molecular biology and bioinformatics. Bioinformatics 2009, 25, 1422–1423.

(76) Michaud-Agrawal, N.; Denning, E. J.; Woolf, T. B.; Beckstein, O. MDAnalysis: A toolkit for the analysis of molecular dynamics simulations. Journal of Computational Chemistry 2011, 32, 2319–2327.

(77) Richard J. Gowers; Max Linke; Jonathan Barnoud; Tyler J.E. Reddy; Manuel N. Melo; Sean L. Seyler; Jan Domański; David L. Dotson; Sébastien Buchoux; Ian M. Kenney; Oliver Beckstein MDAnalysis: A Python Package for the Rapid Analysis of Molecular Dynamics Simulations. Proceedings of the 15th Python in Science Conference. 2016; pp 98 – 105.

(78) Trott, O.; Olson, A. J. AutoDock Vina: Improving the speed and accuracy of docking with a new scoring function, efficient optimization, and multithreading. Journal of Computational Chemistry 2010, 31, 455–461.

(79) Goodsell, D. S.; Lauble, H.; Stout, C. D.; Olson, A. J. Automated docking in crystallography: analysis of the substrates of aconitase. Proteins: Structure, Function, and Bioinformatics 1993, 17, 1–10.

(80) Agarwal, R.; Smith, J. C. Speed vs Accuracy: Effect on Ligand Pose Accuracy of Varying Box Size and Exhaustiveness in AutoDock Vina. Molecular Informatics 2023, 42, 2200188.

(81) Mysinger, M. M.; Carchia, M.; Irwin, J. J.; Shoichet, B. K. Directory of useful decoys, enhanced (DUD-E): better ligands and decoys for better benchmarking. Journal of medicinal chemistry 2012, 55, 6582–6594.

(82) Lagarde, N.; Ben Nasr, N.; Jérémie, A.; Guillemain, H.; Laville, V.; Labib, T.; Zagury, J.-F.; Montes, M. NRLiSt BDB, the manually curated nuclear receptors ligands and structures benchmarking database. Journal of medicinal chemistry 2014, 57, 3117– 3125.

(83) Tribello, G. A. et al. PLUMED Tutorials: A collaborative, community-driven learning ecosystem. The Journal of Chemical Physics 2025, 162, 092501.

