## Supplementary Information for "Advancing in silico drug design with Bayesian refinement of AlphaFold models"

#### Supporting Information

Samiran Sen,[1] Samuel E. Hoff,[1] Tatiana I. Morozova,[1,2] Vincent Schnapka,[1] and Massimiliano Bonomi[*,1]

[1] *Institut Pasteur, Université Paris Cité, CNRS UMR 3528, Computational Structural Biology Unit, Paris, France*

[2] *Current address: CNRS, ENS de Lyon, LPENSL, UMR5672, 69342, Lyon cedex 07, France*

*

### Supplementary Figures

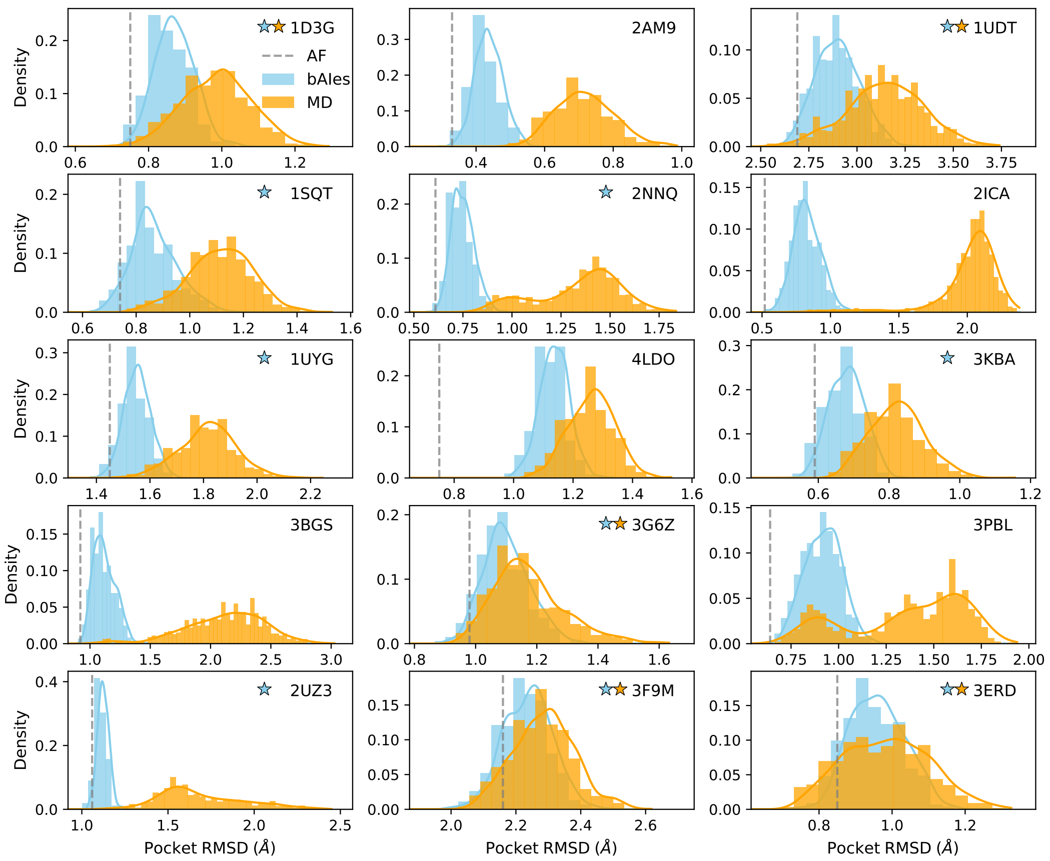


**Supplementary Figure 1: Structural accuracy of ligand binding pockets: MD, bAIes, AF**. Assessment of structural accuracy of ligand binding pockets for the 15 systems in our benchmark set. For each system, the distributions of binding pocket $C_\beta$-RMSD from the reference experimental *holo* conformation are represented for bAIes (blue) and MD (orange) simulations. RMSD of the AF model is represented with a gray dashed line. Orange and blue stars mark whether MD and bAIes, respectively, sample pocket conformations closer to the experimental one than AF. Raw normalized histograms are represented by bars, kernel density estimations as solid lines.

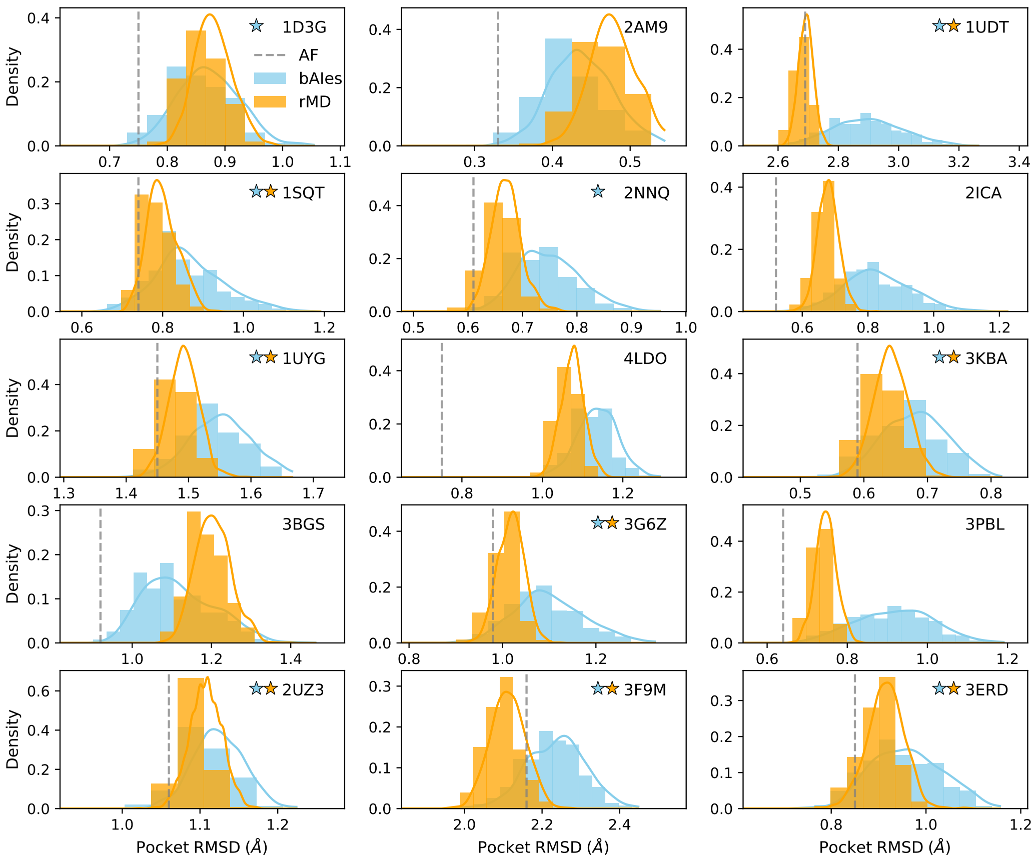


**Supplementary Figure 2: Structural accuracy of ligand binding pockets: rMD, bAIes, AF**. Assessment of structural accuracy of ligand binding pockets for the 15 systems in our benchmark set. For each system, the distributions of binding pocket $C_\beta$-RMSD from the reference experimental *holo* conformation are represented for bAIes (blue) and rMD (orange) simulations. RMSD of the AF model is represented with a gray dashed line. Orange and blue stars mark whether rMD and bAIes, respectively, sample pocket conformations closer to the experimental one than AF. Raw normalized histograms are represented by bars, kernel density estimations as solid lines.

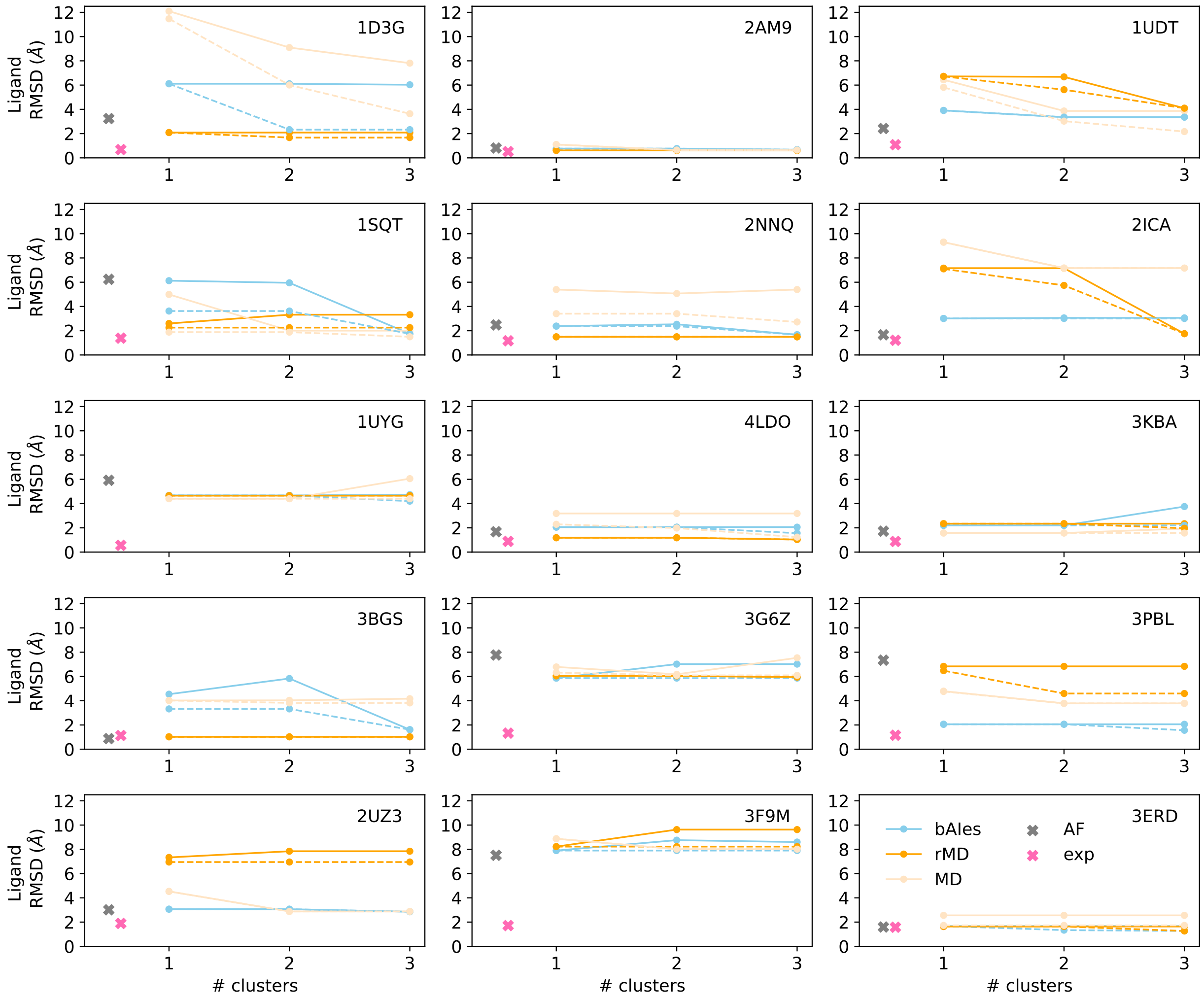


**Supplementary Figure 3: Structural accuracy of native ligand docking poses across clusters**. Lowest ligand RMSDs across all poses calculated cumulatively over the top-3 most-populated clusters (solid lines) using bAIes (blue), rMD (orange), and MD (light orange). Also, the lowest ligand RMSDs among all docked poses (dashed lines) using bAIes (blue), rMD (orange), and MD (light orange). Also, ligand RMSDs for AF (gray, cross) and experiment (pink, cross).

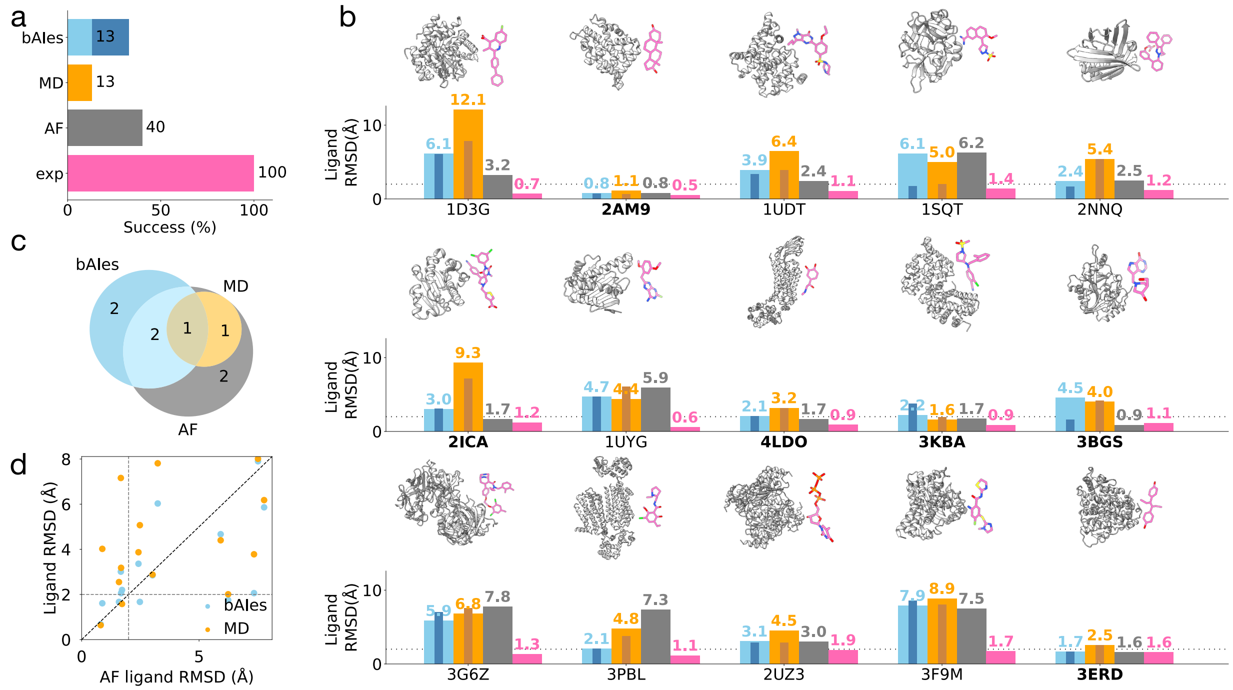


**Supplementary Figure 4: Structural accuracy of native ligand docking poses**. **(a)** Percentage of docking successes across the entire benchmark set of 15 systems using the most populated cluster for bAIes (blue) and MD (orange) as well as using AF (gray) and experimental (pink) structures. Also, the percentage of successes when top three most populated clusters are included for bAIes (dark blue), and MD (dark orange). **(b)** For each system in the benchmark: (*top*) cartoon representation of the experimental *holo* structure, with highlight on the native ligand (pink); (*bottom*) ligand RMSDs with respect to experimental structure obtained from docking against top1 clusters of bAIes and MD, as well as AF, and experimental structures. Bar coloring as in **(a)**. Also, RMSDs when the top3 clusters are included for bAIes (inner dark blue bar) and MD (inner dark orange bar). Systems that resulted in a successful dock by either bAIes, rMD or AF models are marked with the PDB name in bold. **(c)** Diagram illustrating the overlap among successful cases of docking by bAIes, MD, and AF models. Numbers represent unique and shared successes among the models; coloring as in **(a)**. **(d)** Correlation between ligand RMSDs when docking against AF models and either bAIes (blue dots) or MD models (orange dots). The ligand RMSDs values corresponding to the cutoff for successful docking (2Å) are indicated by gray dashed lines.

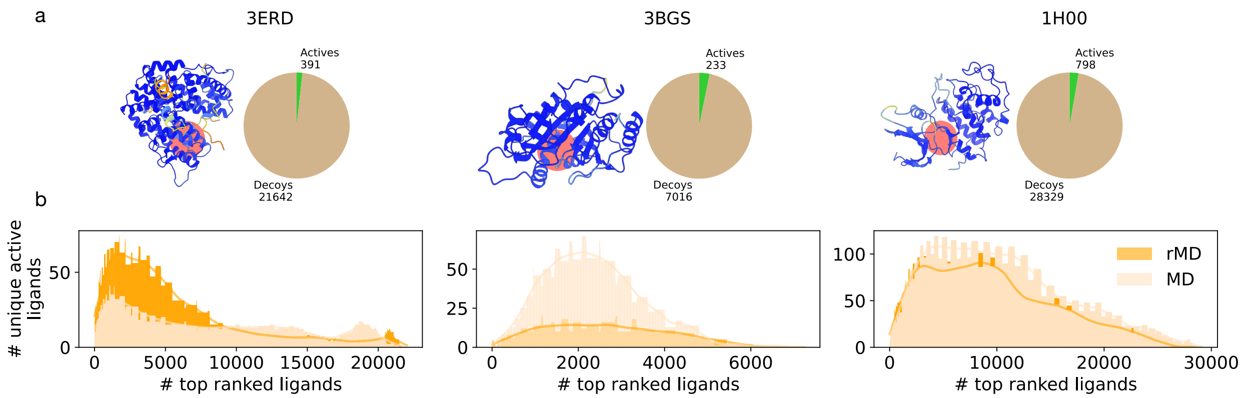


**Supplementary Figure 5: Virtual screening of small molecules libraries.** For 3 selected systems, Estrogen receptor $\alpha$ (PDB 3ERD) (*left*), Purine nucleoside phosphorylase (PDB 3BGS) (*center*), and Cyclin-dependent Kinase (PDB 1H00) (*right*), **(a)** AF model of the protein target (pLDDT-colored), with highlighted binding pocket (coral-colored disc) alongside a pie chart with the number of active (green) and decoy (beige) ligands present in the library. **(b)** Number of unique active ligands identified by unrestrained MD (MD) (light orange) and restrained MD (rMD) (orange) as a function of the number of top-ranked ligands; kernel density estimations (solid lines, colored same as bars).

### Supplementary tables

**Supplementary Table 1: Details of target proteins used in our study**. All structures were obtained by X-ray crystallography. pLDDT is averaged across the residues belonging to the ligand binding pocket.

| Receptor | Receptor ID | PDB | Ligand | pLDDT | Resolution ($\mathring{A}$) |
|---|---|---|---|---|---|
| Dihydroorotate dehydrogenase[1] | PYRD | 1D3G | BRE | 97 | 1.6 |
| Androgen Receptor[2] | ANDR | 2AM9 | TES | 99 | 1.6 |
| Phosphodiesterase 5A[3] | PDE5A | 1UDT | VIA | 95 | 2.3 |
| Urokinase-type plasminogen activator[4] | UROK | 1SQT | UI3 | 98 | 1.9 |
| Fatty-acid-binding protein 4[5] | FABP4 | 2NNQ | T4B | 98 | 1.8 |
| Leukocyte-function associated antigen 1[6] | LFA1 | 2ICA | 2IC | 96 | 1.6 |
| Heat shock protein 90 $\alpha$[7] | HSP90a | 1UYG | PU2 | 96 | 2.0 |
| $\beta_2$ adrenergic receptor[8] | ADRB2 | 4LDO | ALE | 98 | 3.2 |
| Progesterone receptor[9] | PRGR | 3KBA | WOW | 88 | 3.2 |
| Purine nucleoside phosphorylase[10] | PNPH | 3BGS | DIH | 97 | 2.1 |
| Renin[11] | RENI | 3G6Z | A7T | 89 | 2.0 |
| Dopamine $D_3$ receptor[12] | DRD3 | 3PBL | ETQ | 85 | 2.8 |
| Thymidine kinase[13] | KITH | 2UZ3 | TTP | 96 | 2.5 |
| Hexokinase type IV[14] | HXK4 | 3F9M | MRK | 92 | 1.5 |
| Estrogen receptor $\alpha$[15] | ESR1 | 3ERD | DES | 98 | 2.0 |
| Cyclin-dependent Kinase[16] | CDK2 | 1H00 | FAP | 96 | 1.6 |

**Supplementary Table 2: Binding pocket structural accuracy.** Binding pocket RMSD of representatives of the MD top 3 clusters with respect to the reference (*ref.*) as well as closest (*best*) experimental *holo* structures for the 3 systems used in the virtual screening.

| | MD top 3 clusters | | |
|---|---|---|---|
| System | RMSD ref. holo [Å] | PDB best holo | RMSD best holo [Å] |
| Estrogen receptor $\alpha$ | 0.91 | 1GWR | 0.65 |
| | 0.82 | 1GWR | 0.74 |
| | 0.98 | 1GWR | 0.77 |
| Purine nucleoside phosphorylase | 2.47 | 1PF7 | 1.92 |
| | 2.10 | 2Q70 | 1.74 |
| | 2.32 | 2Q70 | 2.20 |
| Cyclin-dependent kinase | 5.48 | 2VTN | 0.95 |
| | 5.43 | 1WCC | 0.90 |
| | 5.41 | 1WCC | 1.16 |

**Supplementary Table 3: Native ligand docking pose accuracy** Ligand RMSD obtained by docking the reference native ligand to their corresponding experimental structures (exp), AF predicted model (AF), and top3 cluster representatives of the rMD and bAIes simulations for the 3 systems used in the virtual screening.

| | | | Ligand RMSD [Å] | | | |
|---|---|---|---|---|---|---|
| System | ref PDB | Ligand | exp | AF | MD | bAIes |
| Estrogen receptor $\alpha$ | 3ERD | DES | 1.58 | 1.59 | 1.64 | 1.67 |
| Purine nucleoside phosphorylase | 3BGS | DIH | 1.13 | 0.88 | 1.02 | 1.61 |
| Cyclin-dependent kinase | 1H00 | FAP | 1.86 | 5.48 | 5.54 | 4.78 |

### References


(1) Liu, S.; Neidhardt, E. A.; Grossman, T. H.; Ocain, T.; Clardy, J. Structures of human dihydroorotate dehydrogenase in complex with antiproliferative agents. *Structure* **2000**, *8*, 25–33.

(2) Pereira de Jésus-Tran, K.; Côté, P.-L.; Cantin, L.; Blanchet, J.; Labrie, F.; Breton, R. Comparison of crystal structures of human androgen receptor ligand-binding domain complexed with various agonists reveals molecular determinants responsible for binding affinity. *Protein Science* **2006**, *15*, 987–999.

(3) Sung, B.-J. et al. Structure of the catalytic domain of human phosphodiesterase 5 with bound drug molecules. *Nature* **2003**, *425*, 98–102.

(4) Wendt, M. D.; Geyer, A.; McClellan, W. J.; Rockway, T. W.; Weitzberg, M.; Zhao, X.; Mantei, R.; Stewart, K.; Nienaber, V.; Klinghofer, V.; Giranda, V. L. Interaction with the S1$\beta$ -pocket of urokinase: 8-heterocycle substituted and 6,8-disubstituted 2-naphthamidine urokinase inhibitors. *Bioorganic & Medicinal Chemistry Letters* **2004**, *14*, 3063–3068.

(5) Sulsky, R. et al. Potent and selective biphenyl azole inhibitors of adipocyte fatty acid binding protein (aFABP). *Bioorganic & Medicinal Chemistry Letters* **2007**, *17*, 3511–3515.

(6) Potin, D. et al. Discovery and Development of 5-[(5S,9R)-9- (4-Cyanophenyl)-3-(3,5-dichlorophenyl)-1-methyl-2,4-dioxo-1,3,7-triazaspiro[4.4]non-7-yl-methyl]-3-thiophenecarboxylic Acid (BMS-587101)A Small Molecule Antagonist of Leukocyte Function Associated Antigen-1. *Journal of Medicinal Chemistry* **2006**, *49*, 6946–6949.

(7) Wright, L. et al. Structure-Activity Relationships in Purine-Based Inhibitor Binding to HSP90 Isoforms. *Chemistry & Biology* **2004**, *11*, 775–785.

(8) Ring, A. M.; Manglik, A.; Kruse, A. C.; Enos, M. D.; Weis, W. I.; Garcia, K. C.; Kobilka, B. K. Adrenaline-activated structure of $\beta$2-adrenoceptor stabilized by an engineered nanobody. *Nature* **2013**, *502*, 575–579.

(9) Kallander, L. S. et al. Improving the developability profile of pyrrolidine progesterone receptor partial agonists. *Bioorganic & Medicinal Chemistry Letters* **2010**, *20*, 371–374.

(10) Rinaldo-Matthis, A.; Murkin, A. S.; Ramagopal, U. A.; Clinch, K.; Mee, S. P. H.; Evans, G. B.; Tyler, P. C.; Furneaux, R. H.; Almo, S. C.; Schramm, V. L. l-Enantiomers of Transition State Analogue Inhibitors Bound to Human Purine Nucleoside Phosphorylase. *Journal of the American Chemical Society* **2008**, *130*, 842–844.

(11) Bezencon, O. et al. Design and Preparation of Potent, Nonpeptidic, Bioavailable Renin Inhibitors. *Journal of Medicinal Chemistry* **2009**, *52*, 3689–3702.

(12) Chien, E. Y. T.; Liu, W.; Zhao, Q.; Katritch, V.; Han, G. W.; Hanson, M. A.; Shi, L.; Newman, A. H.; Javitch, J. A.; Cherezov, V.; Stevens, R. C. Structure of the Human Dopamine D3 Receptor in Complex with a D2/D3 Selective Antagonist. *Science* **2010**, *330*, 1091–1095.

(13) Welin, M.; Kosinska, U.; Mikkelsen, N.-E.; Carnrot, C.; Zhu, C.; Wang, L.; Eriksson, S.; Munch-Petersen, B.; Eklund, H. Structures of thymidine kinase 1 of human and mycoplasmic origin. *Proceedings of the National Academy of Sciences* **2004**, *101*, 17970–17975.

(14) Petit, P.; Antoine, M.; Ferry, G.; Boutin, J. A.; Lagarde, A.; Gluais, L.; Vincentelli, R.; Vuillard, L. The active conformation of human glucokinase is not altered by allosteric activators. *Acta Crystallographica Section D* **2011**, *67*, 929–935.

(15) Shiau, A. K.; Barstad, D.; Loria, P. M.; Cheng, L.; Kushner, P. J.; Agard, D. A.; Greene, G. L. The Structural Basis of Estrogen Receptor/Coactivator Recognition and the Antagonism of This Interaction by Tamoxifen. *Cell* **1998**, *95*, 927–937.

(16) Beattie, J. F.; Breault, G. A.; Ellston, R. P.; Green, S.; Jewsbury, P. J.; Midgley, C. J.; Naven, R. T.; Minshull, C. A.; Pauptit, R. A.; Tucker, J. A.; Pease, J. Cyclin-dependent kinase 4 inhibitors as a treatment for cancer. Part 1: identification and optimisation of substituted 4,6-Bis anilino pyrimidines. *Bioorganic & Medicinal Chemistry Letters* **2003**, *13*, 2955–2960.
